# Predicting Pacific cod spawning habitat in a changing climate

**DOI:** 10.1101/2022.10.04.510851

**Authors:** Jennifer S. Bigman, Benjamin J. Laurel, Kelly Kearney, Albert J. Hermann, Wei Cheng, Kirstin K. Holsman, Lauren A. Rogers

## Abstract

Warming temperatures elicit shifts in habitat use and geographic distributions of fishes, with uneven effects across life stages. Spawners and embryos are particularly sensitive to environmental conditions, with direct impacts of temperature on spawning habitat, as well as indirect connections between their population dynamics and fisheries effort, productivity, and management. Here, we ask how changing environmental conditions and thermal sensitivities of developing embryos confer spatiotemporal variability of thermally-suitable spawning habitat for Pacific cod in the eastern Bering Sea. Specifically, we use bottom temperature values from regionally downscaled global climate models coupled with an experimentally-derived relationship between hatch success and temperature to predict how the extent, mean latitude, and consistency of suitable spawning habitat has changed in the past and may change into the future. We then validate our predictions of suitable spawning habitat with distributions of adults and larvae and examine whether thermal habitat availability relates to recruitment success into the adult cod into the population. We find that the extent and mean latitude of suitable spawning habitat increase over time, particularly if no climate change mitigation occurs in the future. Hotspots of suitable spawning habitat are consistent across shorter time periods but do shift across the Bering Sea shelf by the end of the century. Finally, we find no correlation between the availability of suitable spawning habitat and annual estimates of recruitment. Collectively, our results suggest that as temperatures warm, the availability of suitable spawning habitat will increase and expand spatially and, thus, is not likely to limit recruitment. This work highlights the importance of coupling experimental data with climate models to identify the complex and mechanistic dynamics among temperature, life histories, and ecology, and offers a pathway for examining life stage-specific changes in habitat use and distribution with continued climate change.

## Introduction

Changing environmental conditions have elicited changes in the ecology, physiology, and life histories of marine fishes (Edwards & Richardson, 2004; Parmesan & Yohe, 2003; Poloczanska et al., 2013). One of the most notable, widespread responses to environmental change are shifts in patterns of habitat use and geographic distributions (Burrows et al., 2014; Perry et al., 2005; Pinsky et al., 2013). These include changes in seasonal use of habitat, poleward distributional shifts, and movement of species into deeper waters (Dulvy et al., 2008; English et al., 2021; Fossheim et al., 2015). Although such distributional shifts are likely related to a host of factors, few are as relevant across species and systems as temperature. Changes in temperature have been linked to changes in habitat use and distribution across scales, ranging from individuals tracking preferred temperatures to the large-scale, global redistribution of population and species (Fredston et al., 2021; Marsh & Mueter, 2020; Pörtner & Farrell, 2008). Indeed, thermal limits and tolerance ranges – the range of temperatures a species can physiologically withstand – are thought to shape species distributions and set range boundaries now and in the future (Pörtner, 2021; Sunday et al., 2019; Sunday et al., 2011).

The effect of temperature on organismal physiology is not static across the lifespan of an individual, and thus, thermal limits and tolerance typically change with size and ontogeny (Brown et al., 2016; Dahlke et al., 2020; McKenzie et al., 2020). These changes in thermal sensitivities translate into uneven shifts in habitat use and distribution across life stages (Barbeaux & Hollowed, 2018; Cote et al., 2021; Drost et al., 2016). In particular, developing embryos and spawners are thought to exhibit the narrowest thermal tolerances, resulting in a tight relationship between spawning habitat and temperature (Dahlke et al., 2018, 2020; Pörtner, 2021). For example, reductions in the amount of suitable thermal habitat due to warming disproportionately affects Nassau Grouper (*Epinephelus striatus*) spawners, whose thermal range is narrower than that of nonspawners (Asch & Erisman, 2018). Additionally, compared to other life stages, eggs typically occupy the least amount of habitat area (lowest habitat extent) and have the highest habitat consistency (same locations) over time (Ciannelli et al., 2015, 2021). In turn, these narrow thermal preferences of spawners and embryos collectively result in spatial or temporal (or both) constraints on spawning habitat, which encompasses the habitat of both eggs and spawners (Ciannelli et al., 2021; Rijnsdorp et al., 2009). Thus, spawning habitat has been suggested to not only underlie reproductive potential, but also may represent a limit on species’ ability to adapt to changing environmental conditions (Ciannelli et al., 2021; Dahlke et al., 2020; Porter & Ciannelli, 2018). Spawning habitat may even act as a ‘thermal bottleneck’ that determines species’ vulnerability to a changing climate (Asch & Erisman, 2018; Ciannelli et al., 2021; Dahlke et al., 2020).

Temperature-driven distributional shifts are linked to changes in population dynamics and fisheries productivity, ecosystem dynamics (e.g., species interactions), and socioeconomic aspects such as increased fishing effort and altered management practices (Cheung et al., 2009; García Molinos et al., 2016; Pinsky, Rogers, et al., 2020; Rogers et al., 2019). These factors may impact spawning habitat by influencing the locations of either adult spawners or developing embryos. For example, a reduction in the area or temporal window for successful spawning is correlated with reduced recruitment and stock reproductive potential, as well as altered stock-recruitment dynamics (Dodson et al., 2019; Hayes et al., 1996; Laurel & Rogers, 2020). Additionally, changes in spawning habitat will likely affect the location and movement of adult fish, which in turn can determine catchability in fisheries monitoring surveys, as well as fishing opportunities, as fishers often have limitations on where, when, and how they can fish (Blanchard et al., 2008; Haynie & Pfeiffer, 2013; Rogers et al., 2019). Thus, quantifying how spawning habitat changes with temperature, and whether this confers changes in population and ecosystem dynamics, can help us understand, predict, and mitigate the effects of climate change on fisheries productivity, as well as the livelihoods and economies fishing supports.

The Bering Sea is undergoing rapid environmental change yet supports some of the largest fisheries in the world (Fissel et al., 2019; Grebmeier, 2006; Stabeno & Bell, 2019). In particular, the dynamics of sea ice play a major role in this system, as the timing and extent of seasonal ice affect many other physical and biological factors. A region of cold (<2°C) bottom water on the Bering Sea shelf – the cold pool – is formed during the freezing and melting of seasonal sea ice, and years in which there is a high areal ice extent are associated with a large cold pool (Stabeno et al., 2001; Stabeno, Farley Jr., et al., 2012; Stabeno, Kachel, et al., 2012). The cold pool is thought to act as a barrier between arctic and subarctic communities and drive patterns of distribution and habitat use of fishes in the region (Kotwicki & Lauth, 2013; Mueter & Litzow, 2008; Stevenson & Lauth, 2019). For example, many species avoid the cold pool, and thus shift their seasonal distributions depending on its location and extent (Grebmeier, 2006; Mueter & Litzow, 2008; Stevenson & Lauth, 2019). Recent years have seen a reduction in the extent and duration of sea ice and a concomitant shrinking (or disappearance) of the cold pool, which is thought to underlie the recent northward shifts of many species in the region (Mueter & Litzow, 2008; Stabeno & Bell, 2019; Stevenson & Lauth, 2019). However, it remains unclear whether individuals and species’ responses to changes in temperature are transitory or where redistributions are occurring on longer time scales.

Pacific cod is one of the economically important fishery species that has been observed to expand their summer distribution from the southeastern Bering Sea into the northern Bering Sea following a large and protracted marine heatwave and associated loss of sea ice in the region (Spies et al., 2019; Stevenson & Lauth, 2019). Pacific cod spawn in the winter in the eastern Bering Sea along the outer shelf break and along the Aleutian Islands, particularly around Unimak Island (Neidetcher et al., 2014; Rand et al., 2014; Stark, 2007). Although Pacific cod (like many groundfish species in the region) have been observed in the northern Bering Sea seasonally in summer monitoring surveys, it is unknown whether this area is currently or will become suitable for spawning. Compared to other gadids (e.g., Polar cod *Boreogadus saida*, Saffron cod *Eleginus gracilis*, and Walleye pollock *Gadus chalcogrammus*), the spawning dynamics of Pacific cod are particularly sensitive to temperature as embryos have a narrow thermal range for successful development and hatching (Bian et al., 2014; Cote et al., 2021; Dahlke et al., 2018; Laurel & Rogers, 2020). Pacific cod lay demersal eggs that adhere to the bottom for the duration of development, and thus, spawning dynamics are closely tied to bottom temperature (Alderdice & Forrester, 1971; Laurel et al., 2008). Recent work in the Gulf of Alaska has shown that this narrow thermal tolerance of egg survival reduced the availability of thermally-suitable spawning habitat, which may have contributed to the stark decline in pre-recruit abundance and stock reproductive potential for this species following the marine heatwave in 2014 – 2016 (Laurel & Rogers, 2020). However, in the comparatively colder waters of the Bering Sea, it is unknown whether and when changes in bottom temperature may affect cod populations through potential shifts in thermally-suitable spawning habitat.

Here, we quantify how environmental temperature affects the availability of thermally-suitable spawning habitat (here after, ‘suitable spawning habitat’) across space and time and how this may relate to changes in fisheries productivity for Pacific cod in the eastern Bering Sea. We ask whether (i) the extent or area of suitable spawning habitat varies across space and time, (ii) the mean latitude of suitable spawning habitat shifts northward over time, (iii) spawning habitat suitability is consistent across space and time, and finally, (iv) spawning habitat suitability is correlated with recruitment. To answer these questions, we use bottom temperature from a regional ocean model combined with an experimentally-derived relationship between hatch success and temperature. These data and temperature time series allow us to predict suitable spawning habitat and associated metrics spanning 1970 - 2099 under two emission scenarios of the Shared Socioeconomic Pathway (SSP126 and SSP585). These metrics of spawning habitat suitability allow us to infer important ecological, evolutionary, and economic implications of shifts in habitat use. We additionally validate our predictions of spawning habitat with distributional data on spawning adults and newly-hatched larvae. Ultimately, understanding how spawning habitat dynamics are shifting over time and space will help identify future issues and concerns regarding Pacific cod in the eastern Bering Sea.

## Methods

The Bering Sea encompasses a latitudinal range of > 1000 km between the Alaska Peninsula and the Bering Strait, transitioning from a subarctic to an arctic ecosystem from south to north. The eastern portion of this marginal sea is characterized by a broad, shallow (< 200 m depth) shelf that stretches more than 500 km from shore; it can be divided into the southeastern and northeastern Bering Sea at around 60°N based on differences in physical, chemical, and biological oceanography (Coachman, 1986; Mueter & Litzow, 2008; Stabeno, Farley Jr., et al., 2012; Stabeno, Kachel, et al., 2012). In particular, the northern Bering Sea has a greater areal extent of sea ice and is generally characterized by colder temperatures. The southern Bering Sea can be further subdivided along isobaths into the inner (coastline to 50 m), middle (50-100 m), and outer shelves (100-180 m) based on the unique oceanographic and bathymetric characteristics of each domain (Cheng et al., 2021; Coachman, 1986; Stabeno, Farley Jr., et al., 2012; Stabeno, Kachel, et al., 2012). Broadly, these regions are characterized by differing seasonal mixing and stratification, which impact physical and biological features such as tidal/wind energy, temperature, salinity, nutrient availability, productivity, and food web dynamics (Coachman, 1986; Springer, 1961; Stabeno et al., 2016). For example, the inner shelf is typically well-mixed year-round whereas the middle shelf is well-mixed in the winter but highly stratified in the summer and thus effectively separated from the inner shelf (Stabeno et al., 2016; Stabeno, Farley Jr., et al., 2012). A predominant feature of the middle shelf is the cold pool, a cold (< 2 °C) mass of water along the seafloor that occasionally extends onto the inner shelf in years with extensive sea ice (Stabeno, Farley Jr., et al., 2012). The outer shelf is known for its high productivity resulting from regional tidal mixing, eddies, and transverse circulation (Mizobata et al., 2006; Springer, 1961). Of note, the traditional spawning grounds of Pacific cod are along the outer shelf edge near Zhemchug Canyon, near the Pribilof Islands, and along the Aleutian Islands, particularly on the north side of Unimak Island, all of which are areas of intensive tidal mixing and flow (Neidetcher et al., 2014; Springer, 1961). Indeed, the two major sources of water flowing onto the eastern Bering Sea shelf are the Bering Slope Current that flows along the shelf edge and water from the North Pacific that flows through Unimak Pass (Stabeno et al., 2016).

In recent years, the Bering Sea has shifted from a system dominated by high interannual variability in sea ice, cold pool extent, and temperature to one now characterized by multi-year periods of ‘warm’ years (with low ice extent, reduced/absent cold pool) or ‘cold’ years (higher ice extent, larger cold pool; Overland et al., 2012; Stabeno et al., 2017; Stabeno & Bell, 2019). This is thought to have contributed to a shift in summer habitat use of many groundfishes in the region, including Pacific cod, which were not historically abundant in the northern Bering Sea (Sample & Wolotira, Jr, 1985; Wolotira, Jr et al., 1977). Pacific cod do exhibit seasonal movement from their summer feeding grounds on the inner and middle shelves of the eastern Bering Sea to their winter spawning grounds along the outer shelf edge and along the Aleutian Islands (Neidetcher et al., 2014; Rand et al., 2014; Shimada & Kimura, 1994). While the location of the summer feeding grounds has shifted with temperature (and cold pool extent), the location of winter-spring spawning habitat may be less flexible due to spatial constraints stemming from the narrow thermal range for successful development and hatching of Pacific cod embryos – between 4 to 6 °C (Bian et al., 2016; Laurel & Rogers, 2020).

### Bottom temperature hindcasts and projections

To simulate changes in suitable spawning habitat, we use a regional ocean model known as Bering10K. The Bering10K model is an instance of the Regional Ocean Modeling System (ROMS), with a domain spanning the Bering Sea and northern Gulf of Alaska and including explicit ocean, sea ice, and biogeochemical components (Cheng et al., 2021; Hermann et al., 2021; Kearney et al., 2020, Pilcher et a. 2019). For this study, we used several different simulations from this model. The first, which we refer to as the hindcast, covers the period of 1970 - 2020, and is driven by surface and boundary conditions from the Climate Forecast System operational analysis (Saha et al., 2014). This is a reanalysis product, so this simulation captures the true interannual and decadal variability of the time period. The remaining simulations downscale long-term forecast simulations from the Coupled Model Intercomparison Project Phase 6 (CMIP6; Cheng et al., 2021; Hermann et al., 2021). The downscaled suite includes simulations from three different parent models: Community Atmospheric Model version 6 (CEMS2-CAM6, hereafter ‘CESM’), Geophysical Fluid Dynamics Laboratory Earth System Model version 4.1 (GFDL-ESM4; hereafter, ‘GFDL’), and MIROC-Earth System Version 2 for Long-term simulations (MIROC_ES2L; hereafter, ‘MIROC’). For each parent model, we downscaled the latter portion of the historical simulation (1985 - 2015) and two different emissions scenarios (2015 - 2099): high (SSP126) and low (SSP585) carbon mitigation scenarios (Cheng et al., 2021; Hermann et al., 2021). These particular parent models and emissions scenarios were chosen to capture as much of the envelope of uncertainty from the larger CMIP6 suite as possible given the computing and time constraints of the regional downscaling process.

From each of these simulations, we extracted weekly-averaged bottom temperature, defined as the mean temperature over the bottom 5m of each grid cell. Data were trimmed to include the season (January – April) and bathymetric range (shallower than 250 m) of known spawning (Neidetcher et al., 2014; Rand et al., 2014; Stark, 2007). We also masked the data to include only regions for which the Bering10K bottom temperature output has been validated against observations collected during the Alaska Fisheries Science Center’s groundfish survey (Ortiz & Greg, 2013; Sigler et al., 2010; Kearney et al., 2021; Figure 1a - c). Finally, we trimmed the projection simulations (CESM, GFDL, MIROC) to 2021 – 2099.

**Figure 1.**
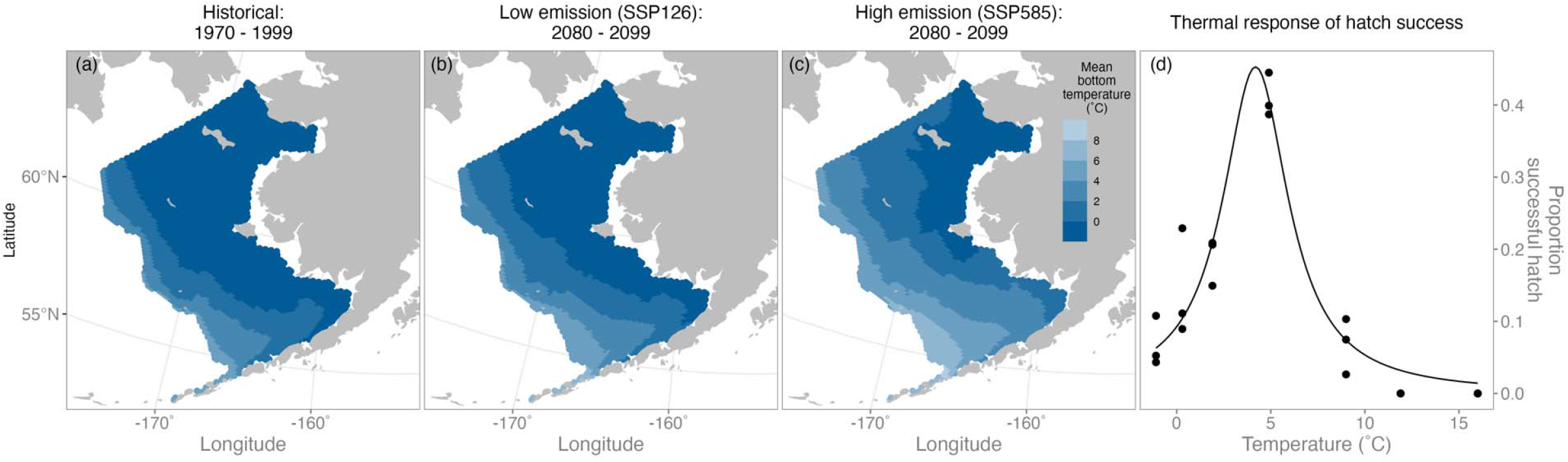
The predicted average bottom temperature in the eastern Bering Sea during March by the end of the century for both emission scenarios is higher than the historical period, and is generally below the optimum temperature for successful hatching of Pacific cod eggs aside from the outer shelf south of ∼57°N. The bottom temperature averaged over a 20-year period in March in the past (a; historical period, 1970 – 1999) and at the end of the century (2080 – 2099) for two emission scenarios (b, low; c, high) from the Bering10K ROMS model. (d) The thermal response of hatch success measured experimentally for Pacific cod as modified from Laurel & Rogers, (2020).

Global climate models – and thus regionally downscaled models driven by global models – can show systematic mismatches between their output and real-world values stemming from a variety of different factors, including coarse resolution, simplified processes, and imperfect understanding of physical and ecosystem processes (Cote et al., 2021; Hawkins et al., 2013). A small bias in simulated bottom temperature could strongly impact our projections of present-day and future habitat suitability due to the narrow, fixed thermal range that defines this habitat. To mitigate these systematic differences, we adjust the projection values using the delta method following Holsman et al., (2020). Essentially this method calculates the bias by comparing the Bering10K results driven by the “free-running” CMIP6 global projections over the reference period, with the corresponding results driven by the observed global conditions over that same time period (the hindcast). The specific equation for adjusting temperature values is:

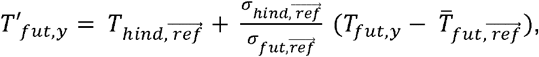

where *T*’_*fut,y*_ is the adjusted projected timeseries, *T*_*fut,y*_ is the raw projected time series, 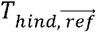 is the mean of the hindcast during the reference years 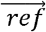 (1980 – 2014), 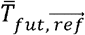 is the mean of the raw projected timeseries during the reference years 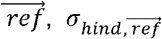 is the standard deviation of the hindcast during the reference years 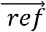, and 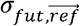 is the standard deviation of the raw projected time series during the reverence years 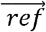. The means and standard deviations of temperature for the reference years were averaged for a given marine sub-region as defined by the Bering Sea Ecosystem Study [BEST] and Bering Sea Integrated Ecosystem Research Program [BSIERP] (see Ortiz & Greg, 2013; Sigler et al., 2010 for more details). We also explored how the choice of reference period affected adjusted temperatures. Specifically, we used two alternative reference periods (1990 – 2014 and 2006 – 2014) but found that the adjusted temperatures were robust to the choice of reference period (Figure S1).

### Spawning habitat suitability and associated metrics

To identify temporal patterns in spawning habitat suitability and provide a basis for calculating area and mean latitude, we calculated an index of spawning habitat suitability. We did so by combining the experimentally-derived relationship between hatch success and temperature from Laurel & Rogers (2020), as described by the Cauchy model (Figure 1d), with the hindcasts and projections of bottom temperature from the Bering10K output. This resulted in a proportion successful hatch per a given grid cell for a given week from January to April for each year spanning 1970 – 2099; these values were averaged on a monthly and yearly basis and were standardized on a scale from zero to one (following Cote et al., 2021; Dahlke et al., 2018). To understand how the area of thermally suitable habitat for spawning changes, we calculated the extent or area of suitable spawning habitat. For this, we summed the area of all grid cells that reached two threshold values – 0.9, termed, “core spawning habitat suitability” and 0.5, termed, “potential spawning habitat suitability” – at any point across the time period examined. This yielded monthly and yearly area estimates for January to April for both core and potential suitable spawning habitat. Third, we calculated the mean latitude of both core and potential spawning habitat area for each year (i.e., for core habitat area, mean latitude would equal the average latitude of all grid cells for which spawning habitat suitability was equal or greater than 0.9) to understand whether core and potential habitat area were shifting northward, as is commonly seen across marine species as temperatures rise (Fossheim et al., 2015). Finally, we examined the consistency of spawning habitat suitability over space and time, which helps us understand how reliable or predictable a particular location is for successful spawning. To do so, we calculated the percent of years a given grid cell was equal to or greater than the two threshold values (0.9 and 0.5, or core and potential suitable spawning habitat).

### Correlating spawning habitat suitability with recruitment

To determine whether the availability of suitable spawning habitat relates to year-class strength (a proxy for recruitment), we obtained estimates of abundance of age-0 Pacific cod, and associated error, from the stock assessment model, an age-structured model fit to survey and fishery data (Thompson et al., 2021). We then estimated the Pearson’s correlation coefficient for the relationship between age-0 abundance (log-transformed) and the annual index of spawning habitat suitability (averaged across space for each year).

### Validating spawn location

To validate thermal habitat model predictions, we examined the distributions of newly-hatched larvae around the months of known spawning (January – April) in the eastern Bering Sea (Neidetcher et al., 2014; Rand et al., 2014; Shimada & Kimura, 1994). Specifically, we mapped distributions of larval density from ichthyoplankton surveys conducted by NOAA Alaska Fisheries Science Center’s Ecosystems & Fisheries-Oceanography Coordinated Investigations (EcoFOCI) for all years (n = 24; 1979, 1991, 1993-1997, 1999, 2000, 2002, 2003, 2005 - 2017) and stations available. These surveys use paired bongo nets with 333 or 505 µm mesh and sample via oblique tows from the surface to 100m depth (or from 10 m off the bottom for shallower water; Matarese et al., 2003; Smart et al., 2012). We limited these data to the months of April, May, and June as larvae were most frequently caught during these months and included only individuals < 6mm in length (larvae are ∼ 4 mm in length upon hatching and grow ∼ 0.30 – 0.40 mm per day; Laurel et al., 2008; Miller et al., 2016). We additionally mapped survey stations for which small larvae were not caught for purposes of comparison. Additionally, we compare our predictions of spawning habitat based on temperature to the distributions of spawning fish reported by Neidetcher et al. (2014) and recent tagging work on Pacific cod in the Bering Sea (Thompson et al. 2021).

## Results

### Bottom temperature hindcasts and projections

Winter bottom waters on the southeastern Bering Sea shelf are predicted to warm considerably by the end of the century (Figure 1a-c, Figure 2a,b, Figure S3). Prior to the beginning of the 21st century, environmental conditions in the southeastern Bering Sea, including bottom temperature, showed high interannual variability. From the early 2000s to the present, the environmental and oceanographic conditions for consecutive years were relatively similar, leading to the categorization of years into “warm” or “cold” stanzas. These stanzas are typically four to six years long and translate into to variable patterns of distribution, abundance, and life history of fishes and lower tropic levels in the region. For example, the species composition and abundance of zooplankton differ between stanzas such that the abundance of large copepods and euphausiids decline in warm years but rebound in cold years. Recruitment of several commercially important species in the region (e.g., Pacific cod, Walleye pollock) is generally lower in warm years compared to cold years (Litzow et al., 2022; Stabeno, Kachel, et al., 2012). In the future, this shift between warm and cold stanzas seems to persist until around the middle of the century, after which temperature increases steadily and no more ‘cold’ years/stanzas are predicted to occur (Figure 2a, b). In contrast to the southeastern Bering Sea, the northern Bering Sea has historically been less thermally variable as it is dominated by a high areal ice extent for almost half of the year, which causes bottom temperatures to remain cold, a trend that persists even into the future (Figure 1b, c, Figure S2, Figure S3). Indeed, the winter bottom temperature for the entire northern Bering Sea remains below 0°C through 2100 under the low emission scenario, and under the high emission scenario, much of the region remains below 0°C with the western edge warming to around 2°C (Figure 1b, c, Figure S3).

**Figure 2.**
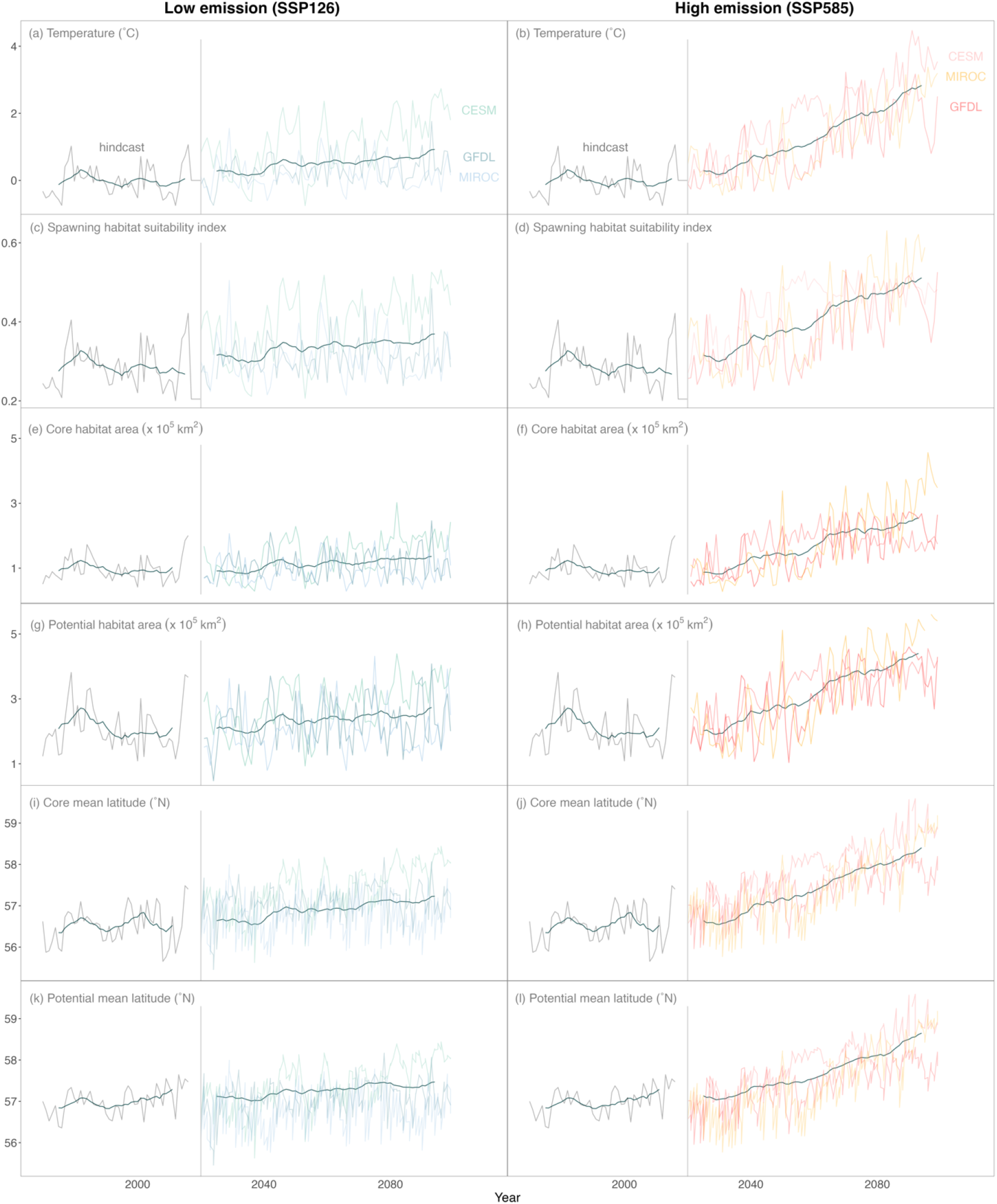
Temperature, habitat suitability, area, and mean latitude are highly variable but increase over time by the end of the century, particularly if no climate mitigation occurs. Time series of (a, b) bottom temperature, (c, d) index of Pacific cod spawning habitat suitability, (e – h) area, and (i – l) mean latitude averaged over January to April of each year for all three global climate models regionally downscaled for the Bering10K ROMS model (colors, see text in panels (a) and (b) for both low (right panels; a, c, e, g, i, k) and high (left panels; b, d, f, h, j, k) emission scenarios. In all panels, the vertical gray line indicates 2020 and the smoothed gray line in the foreground is the 11-year running mean of the hindcast and projection of each respective variable.

Over the 50-year historical period, yearly- and spatially-averaged bottom temperature varied up to 1.9 °C across the eastern Bering sea shelf but did not increase over time (mean slope and 95% Confidence Intervals [CI] = 0.003, −0.005 - 0.012; Figure 2a, b). In contrast, the projected bottom temperature under both emission scenarios did increase over time (mean slope and 95% CI for low [0.009, 0.006 - 0.012] and high [0.034, 0.029 - 0.045] emission scenarios; Figure 2a, b). By mid-century, bottom temperature is predicted to increase up to 2.7°C and 3.8°C under the low and high emission scenario, respectively, and by the end of century, bottom temperature is predicted to increase up to 3.3°C and 4.7°C under the low and high emission scenario, respectively. Under both emission scenarios, the CESM model predicts the greatest temperature increases while the other two models show more modest increases (Figure 2a, b). Such model discrepancy has been documented; for example, Cheng et al., (2021) show that the CESM model predicts temperatures that are warmer than observations while the GFDL and MIROC models predict temperatures that are colder than observations. These differences in temperature projections are due in part to differences in the extent of seasonal sea ice among models (Cheng et al., 2021).

Spatially, the (yearly-averaged) bottom temperature on the outer shelf edge, particularly the southern shelf edge, remained warmer (∼ 4°C) than the middle or inner shelf during the historical period. Over this time, warmer bottom temperatures expanded from the outer shelf edge to the middle and even inner shelf, primarily in the southeastern areas of both domains (Figure S2). The middle and inner shelves north of 57°N generally remained cooler (< 0°C; Figure S2). In the future, warmer bottom temperatures are projected to spread across the shelf from the outer edge to the inner shelf, but largely only in the southeast, particularly south of 59°N (Figure 1c). Similar to the historical period, the middle and inner shelves remain cooler, even by the end of the century. However, warmer bottom temperatures do expand along the northern margin of the study area (around 65°N) by the end of the century (Figure 1c).

### Spawning habitat suitability and associated metrics

#### Index of spawning habitat suitability

The dynamics of bottom temperature over space and time translate into similar dynamics of spawning habitat suitability for both the historical and future periods (Figure 2c, d). Overall, the spawning habitat suitability index, while highly variable over the entire time period, is projected to increase by the end of the century. Such increases in spawning habitat suitability are due to large regions of the Bering Sea shelf where bottom temperatures do not rise above the optimum temperature for successful hatching (Figure 1d).

From 1970 - 2020, the spawning habitat suitability index varied considerably (Figure 2c, d). As with temperature during this time, the spawning habitat suitability index was more similar in consecutive years until the thermal regime switched (i.e., from cold to warm stanza). However, there was no significant directional trend in spawning habitat suitability over the course of the entire historical period, likely due to the colder bottom temperatures remaining on the eastern Bering sea shelf (mean slope and 95% CI = −0.0003, −0.001 - 0.001). In the future, the spawning habitat suitability index is predicted to rise, particularly under the high emission scenario (mean slope and 95% CI for the low [0.003, 0.002 - 0.004] and high [0.008, 0.0004 - 0.001] emission scenario; Figure 2c, d). Indeed, compared to 1970, the spawning habitat suitability index is predicted to increase by over 110% and 181% under the low and high emission scenario, respectively. As with the projected temperature time series, the CESM model predicts larger increases in the spawning habitat suitability index compared to the GFDL and MIROC models (Figure 2c, d).

Spatially, suitable spawning habitat remained high along the outer shelf edge and expanded across the shelf to the southeast middle shelf during the historical period (Figure 3). In the future, areas of high spawning suitability are predicted to shift slightly away from the outer shelf edge towards the middle and inner shelves, particularly in the southern portion of the outer shelf edge (Figure 3). Indeed, a hotspot for which spawning suitability remained high throughout the historical period – the southern portion of the outer shelf edge south of ∼57°N – became less suitable over time (particularly under both emission scenarios of the CESM model). Finally, the northern Bering Sea is predicted to be unsuitable for spawning across the historical and forecasted time period as it remains colder than optimum temperature for egg development and hatching to the end of the century (Figure 3).

**Figure 3.**
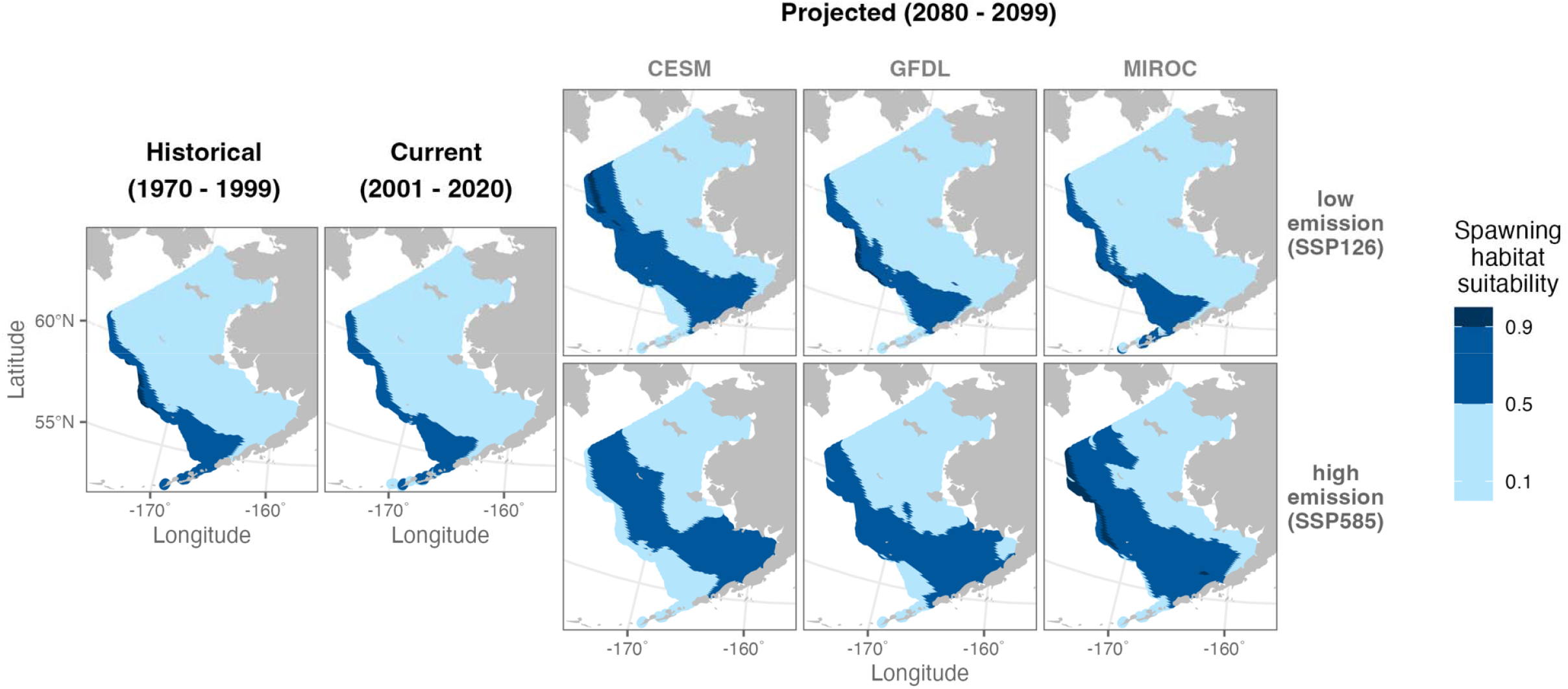
Core and potential suitable spawning habitat are concentrated on the outer shelf and shift and expand spatially towards the middle shelf by the end of the century, particularly under the high emission scenario. Maps of both core (spawning habitat suitability 0.9) and potential (spawning habitat suitability 0.5) suitable spawning habitat for the historical (1970 – 1999), current (2001 – 2020), and last 20 years of the projected period (2080 – 2099; indicated by the black bold text labels above the plots) for Pacific cod. Spawning habitat suitability was averaged across each respective time period and for the last 20 years of the projected period; maps are shown for each global climate model (columns, see text) and emission scenario (rows, see text).

### Area of spawning habitat suitability

Overall, the area of core (spawning habitat suitability 0.9) and potential (spawning habitat suitability 0.5) suitable spawning habitat did not increase over the historical period (mean slope & 95% CI for scaled core [0.002, −0.003 - 0.007] and potential [0.004, −0.005 - 0.006] spawning habitat area). However, the area of both core and potential suitable spawning habitat is projected to increase in the future under the low (mean slope and 95% CI for core [0.003, 0.001 - 0.044] and potential [0.0003, 0.002 - 0.004]) and high (mean slope and 95% CI for core [0.0008, 0.005 - 0.11] and potential [0.0009, 0.006 - 0.012]) emission scenarios (Figure 2e-h). Historical area was variable from 1970 – 2020 and ranged ∼ 200,000 km^2^ (300,000 km^2^) for core (potential) suitable spawning habitat area. By mid-century, core suitable spawning habitat area is projected to increase by ∼40% and ∼80% (i.e., ∼280,000 km^2^ and 360,000 km^2^) under the low and high emission scenario, respectively. Potential suitable spawning habitat area is projected to increase by ∼4% and 41% (i.e., ∼ 311,000 km^2^ and 423,000 km^2^) by mid-century under the low and high emission scenarios, respectively. Both core and potential suitable spawning habitat area continue to increase from mid- to end of the century under the high emission scenario (core area is projected to increase by an additional ∼ 32,000 km^2^ and potential by ∼ 42,000 km^2^; i.e. a 96% and 55% increase over historical area), but under the low emission scenario, both core and potential suitable spawning habitat area are projected to decrease slightly between mid- to late century representing a 30% and −7% change from historical area (core decrease from mid-century of ∼ 20,000 km^2^ and potential by ∼ 32,000 km^2^).

Spatially, the majority of core habitat area during the historical period was concentrated on the outer shelf edge. In contrast, potential area increased and expanded across the shelf during this time, from the outer shelf onto the middle and inner shelves south of ∼ 57°N (Figure 3). During the historical period, the entire northern Bering Sea remained below the threshold of potential habitat (0.5) due to the temperature remaining below 0°C (Figure 3). In the future, suitable spawning habitat area – particularly potential habitat – is projected to increase and expand across the eastern Bering Sea onto the middle and inner shelves by the end of the century, especially under the high emission scenario (Figure 3). For two of the global climate models – CESM and GFDL, both potential and core habitat areas shift slightly inshore, occupying the southern extent of the middle and even inner shelves (south of 57 - 60°N, Figure 3). Similar to the historical time period, the northern Bering Sea is not projected to become thermally suitable for spawning by the end of the century under any model and scenario, as it remains below the range of temperatures that confer a high probability of hatch success (Figure 3).

### Mean latitude of spawning habitat suitability

The mean latitude of both core and potential spawning habitat area shifts northward over time, particularly under the high emission scenario (Figure 2i-l, 4). Compared to the mean latitude for core and potential spawning habitat area averaged across the hindcast period (1970 – 2020), the projected increase in mean latitude by the end of the century is around two degrees latitude for both thresholds, depending on parent model and scenario (Figure 2i-l, 4). Under both emission scenarios, projected increases in mean latitude are roughly similar between core and potential habitat (Figure 2i-l). Finally, the average location of both core and potential spawning habitat – as measured by the mean latitude and longitude – shifts north over time and the spread of locations east to west narrows (Figure 4).

**Figure 4.**
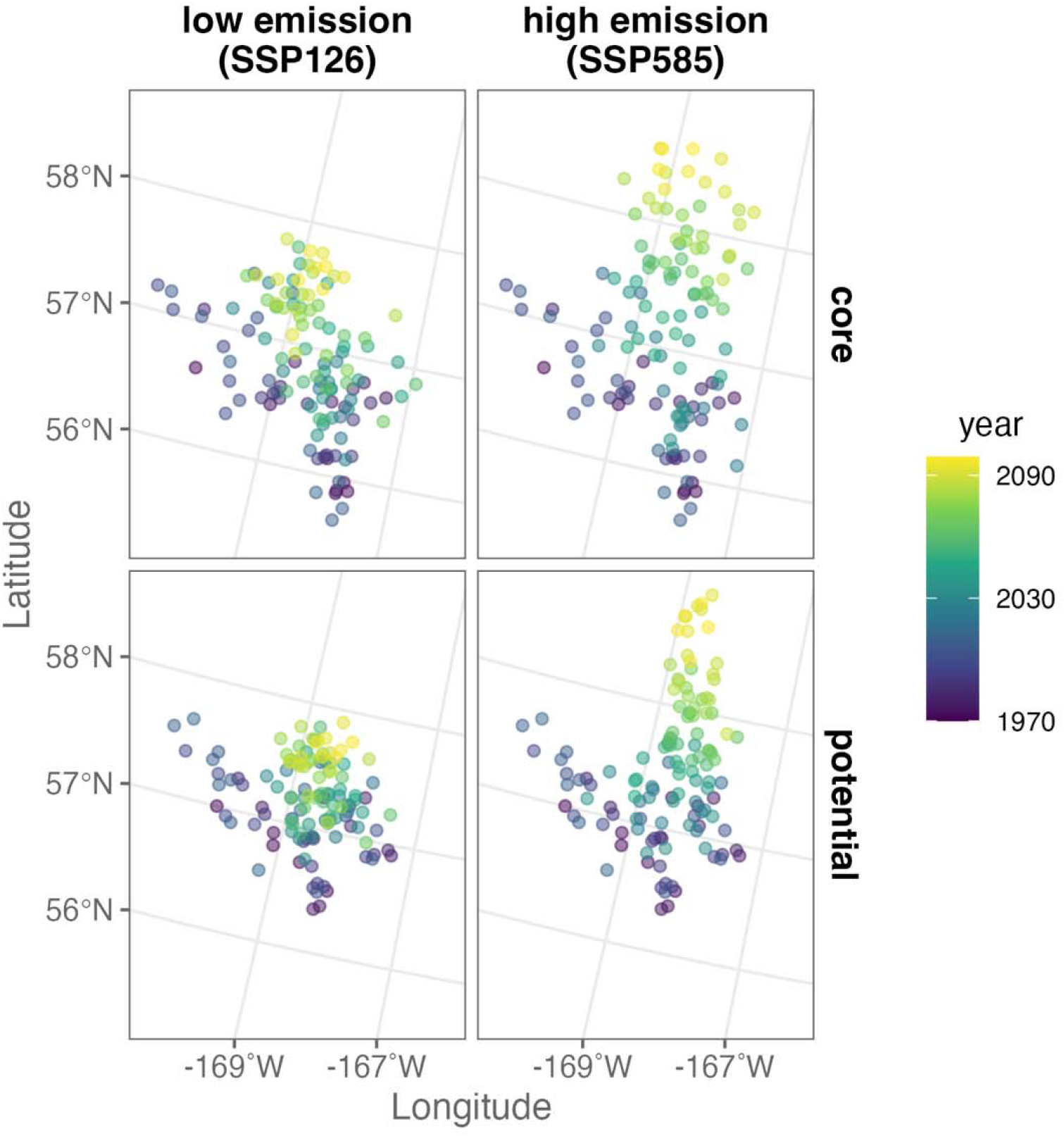
The mean latitude (°N) and longitude (°W) of both core and potential suitable spawning habitat is shifting northward over time and narrowing from east to west. The yearly-averaged mean latitude and longitude of both core (spawning habitat suitability >0.9, top row) and potential (spawning habitat suitability ≥0.5, bottom row) suitable spawning habitat for Pacific cod under low (left column) and high (right column) emission scenarios. For each emission scenario, the projected mean latitude and longitude change is averaged across the three global climate models.

### Consistency of spawning habitat suitability

Spawning habitat suitability values are relatively consistent in a given location across shorter time periods (decades) but shift slightly and expand across the shelf over the course of the entire time frame (130 years), similar to the changes in spawning habitat area (Figure 5). During the historical period, the outer shelf edge was consistently suitable core habitat, as ∼75% or more of years had a spawning habitat suitability index 0.9. For locations just east of the shelf break, the percent of years that were core habitat vary from ∼ 25 – 75% (Figure 5). In the future, areas that are consistently suitable core habitat shift slightly inshore onto the middle shelf over time, particularly south of 57°N. Across this time frame (2021 – 2099), areas that are consistently core habitat vary from having 50 – 100% of years with spawning habitat suitability index values 0.9; higher values are typically found on the northern end of the outer shelf edge and decline toward the southern portion (although the high emission scenario of the MIROC model predicts high consistency on the middle shelf). While both emission scenarios predict similar patterns of consistency for core habitat area, the high emission scenario predicts a further shift towards the inshore region (Figure 5). For potential habitat (spawning habitat suitability index 0.5), areas that are consistently suitable during the historical period remained consistently suitable potential habitat across all years spanning 1970 – 2020). These regions of historic suitability expanded inshore from the outer shelf edge and from the southern portion of the middle shelf onto the southern portion of the inner shelf (Figure 5). This pattern continues in the future, as potential habitat is consistently found on the middle shelf for a high number of years, and under the high emission scenario, the southern region of the inner shelf (Figure 5). Similar to the pattern for core habitat, the outer shelf edge becomes less consistently suitable over time, particularly under both scenarios of the CESM model, as areas that are consistent shift inshore (Figure 5). The northern Bering Sea does not harbor consistently suitable core or potential habitat from 1970 – 2099, with the exception of a small area forecasted by the MIROC model, and to a lesser extent, the CESM model, under the high emission scenario (Figure 5).

**Figure 5.**
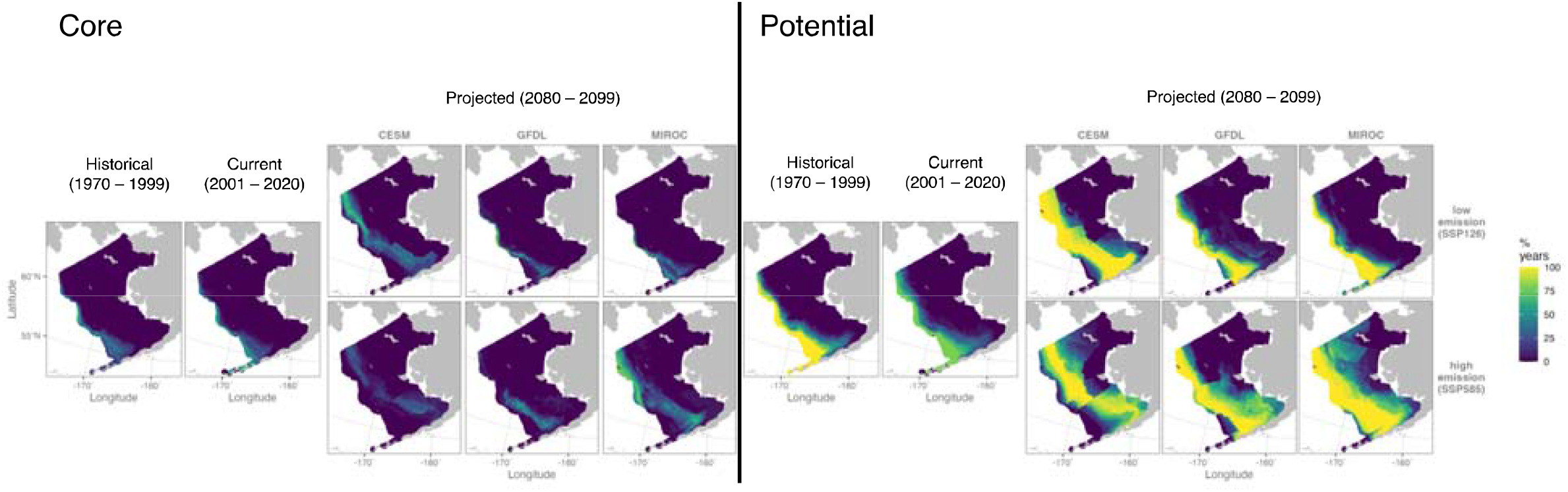
Core and potential suitable spawning habitat are relatively consistent across shorter time periods (decadal scale) but do expand and shift spatially by the end of the century. Maps of the consistency of Pacific cod core (spawning habitat suitability ≥0.9, left panel) and potential (spawning habitat suitability ≥0.5, right panel) suitable spawning habitat for the historical period (1970 – 1999) and current period (2001 – 2020) compared to that projected by the end of the century (2080 – 2099) for all three global climate models (columns of the projected maps) under both low and high emission scenarios (rows of the projected maps). Spawning habitat suitability was averaged across each respective period.

### Spawning habitat suitability and recruitment

Over the time span of years in which abundance estimates of age-0 fish were available (1977 – 2020), we did not find a significant relationship between spawning habitat suitability and abundance (Pearson’s correlation coefficient = −0.25, p = 0.099; Figure 6).

**Figure 6.**
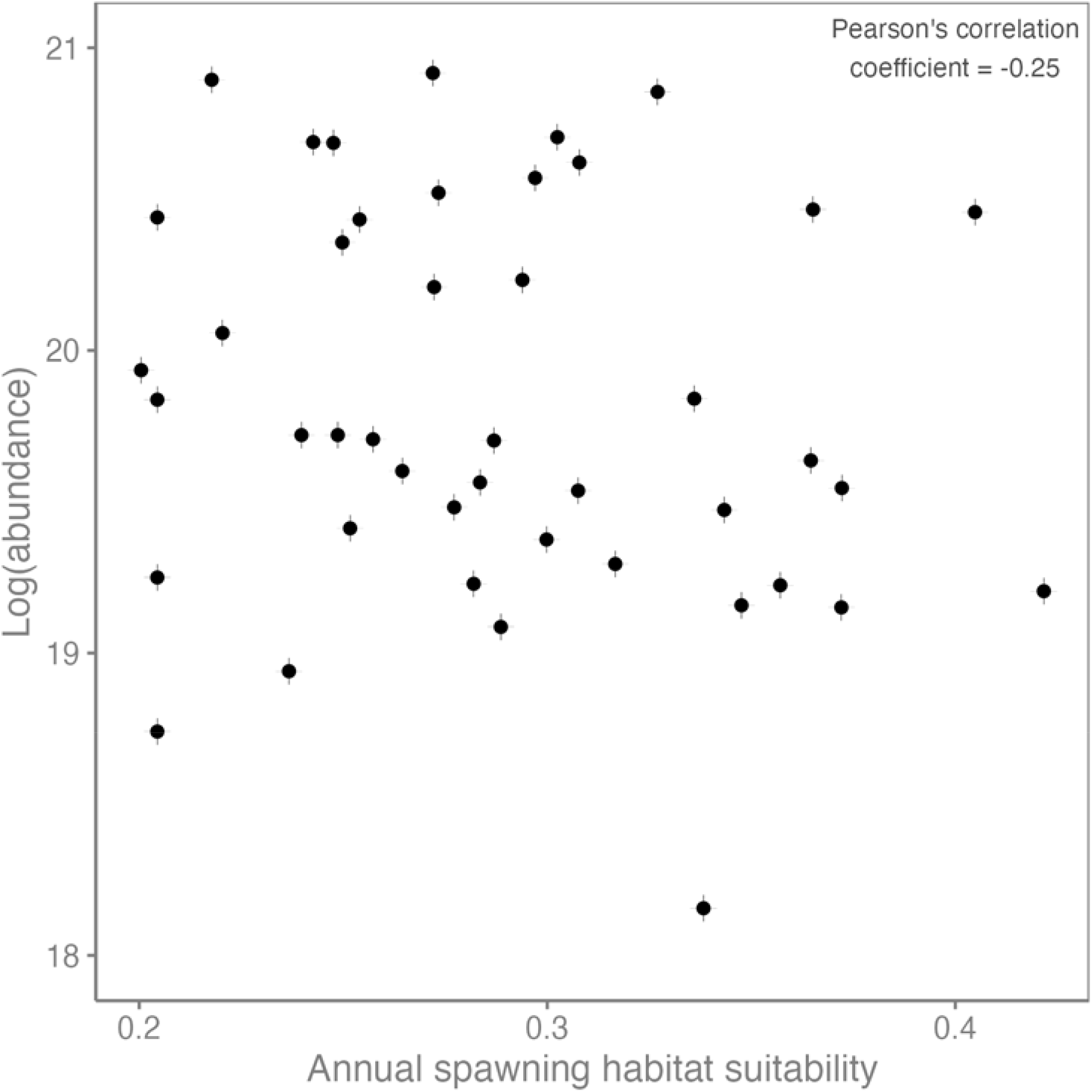
No relationship exists between the annual index of spawning habitat suitability for Pacific cod and the (log) abundance of age-0 fish from 1977 - 2020.

### Validating spawn location

The distributions of both newly-hatched larvae and spawning adults validated our predictions of spawning habitat based on temperature. Specifically, catches of small-sized larvae were concentrated along the outer shelf and along the Aleutian Islands (Figure 7), matching the locations predicted to be sites of spawning habitat based on temperature (Figure 3, “historical” and “current” panels). The distribution of fish in pre-spawning, spawning, and spent stages has typically been concentrated on the outer shelf edge and along the Aleutian Islands, particularly around Unimak Island; additionally, recovery locations of tagged adult Pacific cod during the height of the spawning season (February and March) were concentrated along the outer shelf edge and less so, near Unimak Island (Neidetcher et al. 2014; Thompson et al. 2021).

**Figure 7.**
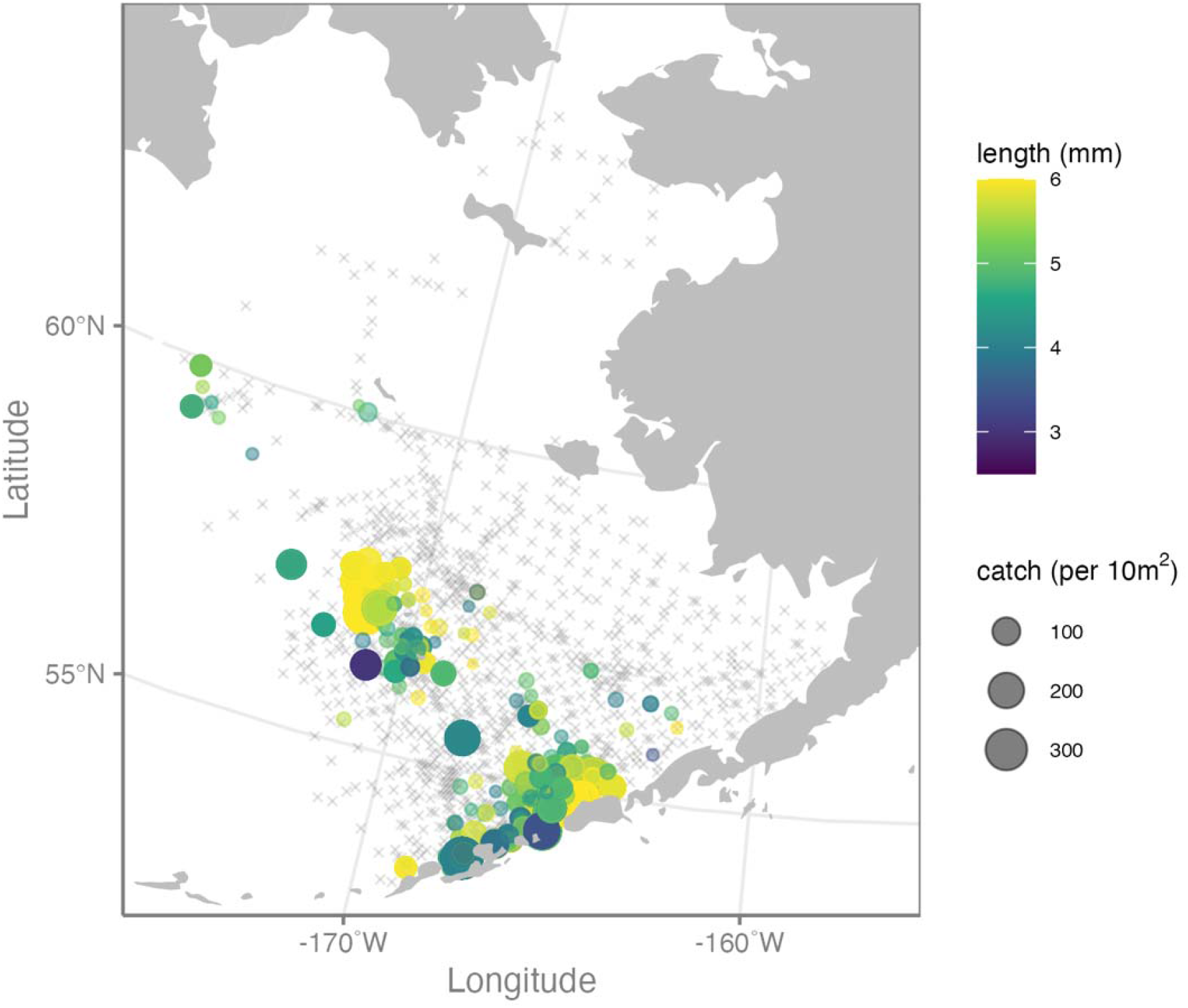
Observations of larval catches validate our predictions of suitable spawning habitat based on temperature. Empirical observations of catch of small Pacific cod larvae (< 6mm) from fisheries surveys conducted in the Bering Sea during April – June for as many years as possible (1979, 1991, 1993-1997, 1999, 2000, 2002, 2003, 2005 – 2017). Also shown (indicated by the black x’s) are the station locations for which small larvae were not caught; these station locations are averaged to the closest latitude and longitude.

## Discussion

Overall, our work suggests that changing thermal conditions in the eastern Bering Sea have affected and will continue to affect the spawning dynamics of Pacific cod. While the availability of thermally-suitable spawning habitat – in terms of a yearly index of spawning habitat suitability (indicating proportion of hatch success/egg survival) and areal extent – is projected to increase by the end of the century, our projections suggest that historical spawning sites will become less thermally suitable over time and suitable spawning habitat will shift slightly northward. Despite the tight mechanistic relationship between temperature and hatch success, which allows us to use bottom temperature to successfully predict the location of spawning, our work suggests that the availability of suitable spawning habitat is not likely constraining regional stock productivity (recruitment). However, the spatial shift in suitable spawning habitat by the end of the century may have other (and possibly unanticipated) consequences. These consequences include physiological costs such as higher metabolic demand from increased temperature or a change in the migration distance from summer feeding areas to spawning sites (sensu Cote et al. 2021). Similarly, spatial shifts in spawning could result in novel species interactions or have socioeconomic implications such as cost and effort for the fishery (Rogers et al., 2019; Sumaila et al., 2011). Although distributional shifts are one of the most well-known responses to warming (Burrows et al., 2014; Pinsky et al., 2013; Pinsky, Selden, et al., 2020), our work highlights how such shifts may be size- and season-dependent, as well as leverages a mechanistic relationship and global climate model output to understand how thermal habitat has varied in the past and forecast how it will vary in the future.

We found an overall increase in the availability of suitable spawning habitat, as indicated by both the increase in the spawning habitat suitability index and areal extent of both core and potential suitable spawning habitat. Expansions of habitat area with warming, just as contractions, have been documented for many species (Bennie et al., 2013; Eaton & Scheller, 1996; Kleisner et al., 2017). Such changes in thermal habitat can be dependent on life stage; for example, by the end of the current century, thermally-suitable habitat for spawning adults of Nassau grouper is projected to decline by 82% compared to 46% for non-spawning adults (Asch & Erisman, 2018). Changes in thermal habitat for earlier life stages, as well as their effects, are less well documented. For gadids specifically, both increases and decreases in suitable spawning habitat with warming (based on experimentally-derived relationships between temperature egg survival as used here) have been found, with differences attributed to the nature of the temperature changes and life histories of the individual species (i.e., subarctic vs. arctic species, shape of the thermal response curve, nature of temperature changes in the region; Cote et al., 2021; Dahlke et al., 2018; Laurel & Rogers, 2020). For Polar cod, an arctic species with egg survival highest in waters < 3°C, spawning habitat suitability is projected to decrease in both the Norwegian Sea (by ∼67% in the warmest areas) and in the Labrador Sea and Northwest Atlantic (by ∼21%) by the end of the century (Cote et al., 2021; Dahlke et al., 2018). Atlantic cod, a subarctic species whose eggs have a broader range of temperatures suitable for survival compared to both Polar and Pacific cod (highest survival between 0° and 9°C), can expect regional increases and decreases in suitable spawning habitat by the end of the century (Cote et al., 2021; Dahlke et al., 2018). Using the same experimentally-derived relationship between temperature and egg survival as used here, Pacific cod spawning habitat suitability is expected to increase in the Labrador Sea by the end of the century (by 19%; Cote et al., 2021). However, decreases in spawning habitat suitability have also been documented for this species following a marine heatwave (Laurel & Rogers 2020). We found rather large increases in spawning habitat suitability and area for Pacific cod in the eastern Bering Sea at mid-century with warming temperature under both emissions scenarios. The increases in temperature on the middle and inner shelves over time facilitate the expansion of suitable spawning habitat onto the shelf and translate into increases in area that are enough to counter the decreases on the outer shelf edge. However, the implications of this increase in spawning habitat suitability and area are less well understood. For example, it is unclear whether Pacific cod will use the increased area that is projected to become available with warming. Indeed, Dahlke et al., (2018) noted that while observed spawning for Atlantic cod and Polar cod occurred entirely within the areas predicted to be thermally suitable, their predictions of area were greater than that where spawning actually occurred. While we know less about the area of spawning habitat for Pacific cod in the eastern Bering Sea, the general locations of known spawning are also within the areas predicted to be thermally suitable. However, in our case, it seems that the area currently predicted to be thermally suitable for Pacific cod is not larger than the current areas of known spawning. Thus, Pacific cod spawning activity will largely depend on movement and behavior across these emerging thermal habitats (Ciannelli et al., 2015; Cote et al., 2021; Dahlke et al., 2018).

Despite our finding of an overall increase in the availability of suitable spawning habitat, we also found that sites historically suitable for spawning are not projected to be so by the end of the century. Specifically, the outer shelf edge – a hot spot for current and past spawning– is projected to become warmer than is optimal by the middle of the century for successful development and hatching of eggs (Figure S3; Laurel & Rogers, 2020). Because spawning habitat is thought to represent a confluence of ideal conditions for multiple life stages (spawning adults, larvae, eggs) in addition to temperature (e.g., prey availability, refugia form predators, connectivity between spawning sites and nursery grounds), a shift in spawning location may impact the survival and transport of newly-hatched larvae (Agostini & Bakun, 2002; Ciannelli et al., 2015; Claydon, 2004). If spawning locations shift spatially (or temporally) based on the physiological limits of eggs, new habitats (or the same habitat at a different time) may not offer the necessary conditions that offer the other requirements and characteristics of spawning habitat. One possible advantage to the predictions of increased thermal spawning habitat on the middle and inner shelves is the potential for overlap with habitat identified as important for juvenile survival (Farley et al., 2016; Hurst et al., 2012; Miller et al., 2016). Of course, this assumes that juvenile habitat quality will remain high in these middle and inner shelf regions, which will need to be elucidated with further work. Finally, shifts in spawning aggregations have the potential to affect the logistics and operational costs of fisheries that target such aggregations (Asch & Erisman, 2018; Dahlke et al., 2018; Pinsky & Fogarty, 2012). In the Bering Sea, for example, the fishery for Pacific cod occurs in the winter and is centered on the outer shelf edge and along the Aleutians, sites known to be historically important for spawning adults (Neidetcher et al., 2014; Rand et al., 2014).

As with many studies examining changes in geographic distributions and ranges, we found an overall northward shift in terms of the mean latitude of suitable spawning habitat (Fossheim et al., 2015; Kleisner et al., 2017; Pinsky et al., 2013). However, we note that this northward shift does not include the northern Bering Sea, a region that has become important to both juvenile and adult Pacific cod (and other groundfish species) in the summer months (Stevenson & Lauth, 2019). The warmer than optimal temperatures for spawning in the northern Bering Sea in the winter suggest that Pacific cod, if they continue to move to the northern Bering Sea in the summer, must migrate back to the outer shelf and Aleutian Islands for spawning in the near future and towards the mid and end of the century, the middle and inner regions of the eastern Bering Sea shelf. While changes in habitat use and distributions have been well-documented for later life stages, many large-scale studies examining range shifts across species do so for adults or combine multiple life stages (e.g., juveniles and adults; (Cheung et al., 2013; Morley et al., 2018; Perry et al., 2005). Our study highlights a shift in biogeography from an early life history perspective.

The patterns of ‘warm’ and ‘cold’ stanzas in the recent past of the Bering Sea facilitated a relatively large area of habitat that was consistently thermally suitable (in terms of the % years) for spawning within a decade. However, the locations of consistent habitat did shift over longer time periods as the middle and inner regions of the shelf became more thermally suitable over time, and the outer shelf edge, particularly the southern portion, became less thermally suitability over time. Based on current knowledge, Pacific cod spawn in the same locations from year to year (Neidetcher et al., 2014; Rand et al., 2014). Such patterns indicate that spawning habitat and early life stages of this species are spatially constrained, meaning that they are limited to specific locations that provide the conditions and characteristics for spawning and larval survival (Beck et al., 2001; Ciannelli et al., 2015, 2021). Indeed, for a species to successfully reproduce, the conditions and characteristics of spawning habitat needed to maximize offspring survival must be met over the long-term, and not fluctuate greatly from year to year (Ciannelli et al., 2021). Our finding that the locations of consistent thermally-suitability habitat are similar for shorter periods of time but do shift over the course of the century may indicate reduced reproductive success in the future. For example, Pacific cod spawning habitat seems to be spatially constrained, but warming temperatures may cause it to shift geographically. Alternatively, reproductive success may be less affected if the timing of spawning (spawning phenology) shifts, as is found in other species (Rogers & Dougherty, 2019). However, we were unable to assess this here due to a lack of detailed data on the timing of reproduction of Pacific cod in the Bering Sea.

We did not find a strong relationship between recruitment and spawning habitat suitability. This finding is likely due to the fact that the availability of suitable spawning habitat – based on our predictions from temperature – is not limiting the reproductive potential of Pacific cod in the eastern Bering Sea. Other work on assessing the link between suitable spawning habitat availability and population dynamics has found a relationship between the two, but this is likely due to the find of a reduction, and not increase in habitat that is suitable for spawning (Dodson et al., 2019; Laurel & Rogers, 2020). For example, Laurel and Rogers (2020) found that a reduction of spawning habitat suitability for Pacific cod following a heatwave in the Gulf of Alaska, and thus poor hatch success, was linked to lower abundance of larvae, age-0 juveniles, and age-3 recruits. Likewise, Dodson et al. (2018) revealed that larval abundance of herring (*Clupea harengus*) was higher when water temperatures were optimal with respect to hatch success over a 23-year time series. While not based on thermal habitat specifically, a loss of settlement habitat due to warming for some coral reef fishes – habitat supporting survival of early life stages – is thought to have contributed to declines in abundance (Munday et al., 2008). The relationship between thermal spawning habitat availability and recruitment or productivity may even be nonlinear; for example, Laurel et al. (*in review*) found a threshold response such that recruitment failure was only evident when thermal spawning habitat metrics fell below a certain value. To our knowledge, studies that have found increases in spawning habitat suitability, as we found here (e.g., Cote et al., 2021; Dahlke et al., 2018) have not assessed whether this increase is related to recruitment or population dynamics. In the case of Pacific cod in the eastern Bering Sea, it appears that other factors, and not the availability of suitable spawning habitat, are related to recruitment and stock dynamics (Farley et al., 2016; Miller et al., 2016). Indeed, recruitment of Pacific cod in the Bering Sea, and other groundfish species, is related to a host of factors including climactic variability (e.g., sea ice, winds) and diet and species interactions (predation, prey availability) to name a few (Hollowed et al., 2001; Houde, 2008; Litzow et al., 2022; Rogers et al., 2021). Thus, while warming is not affecting Pacific cod recruitment as mediated by spawning habitat dynamics, warming is undoubtedly affecting Pacific cod and other species in other, and possibly unanticipated, ways. While beyond the scope of our study, future work might consider incorporation of our suitable spawning area index as part of a comprehensive climate-enhanced stock assessment model, which could evaluate the relative role of spawning habitat to recruitment and population productivity.

Although predicting distributions and habitat use from temperature alone has shown to be quite successful across regions, species, and time frames, ecological characteristics such as distributions are complex and are the result of many interactive factors (Pinsky et al., 2013; Pörtner & Farrell, 2008; Sunday et al., 2011). While we note that our predictions of spawning habitat based on temperature alone do match observations of known spawning location based on tagging data (Bryan et al., 2021; Thompson et al., 2021), as well as the winter distributions of spawning adults and newly-hatched larvae, our predictions do rely on several assumptions. First, undoubtedly, temperature is not the only factor governing species distributional patterns and, thus, our predictions of spawning habitat based on temperature preferences are likely a first approximation as to realized patterns of spawning habitat in the past, present, and future. It is likely that including other factors, e.g., oxygen, prey availability, and sediment information, would be future steps to refine our predictions of thermal habitat. As relationships between hatch success and other factors, particularly oxygen, become available, future work can include such relationships. Second, there is uncertainty in the projections of temperature from global climate models. While the Bering10k model temperature hindcasts and projections are validated against past observations and show great skill in predicting abiotic factors, forecasting future environmental conditions is nontrivial (Cheng et al., 2021; Hermann et al., 2021; Kearney et al., 2020). Thus, incorporating other factors into explanatory and predictive models of species’ shifting distributions, such as those mentioned earlier (oxygen, sediment data), would not only build on the temperature relationships used here but also likely reduce the uncertainty in predictions of distributions based on temperature hindcasts and forecasts. Third, it is common that species shift to deeper water with warming (Baudron et al., 2019; Dulvy et al., 2008). Although Pacific cod may respond to warming by moving to colder, deeper waters, eggs are demersal and thus, the location of spawning will likely shift latitudinally and longitudinally, as eggs are already on the seafloor (Alderdice & Forrester, 1971; Laurel et al., 2008; Laurel & Rogers, 2020). Fourth, the use of a static thermal response curve of egg survival across time assumes no plastic or evolutionary change in thermal preferences. While it is likely that thermal preferences are plastic and adaptable (although recent work suggests that across species, upper thermal limits change < 0.1°C over a million years; Comte & Olden, 2017), understanding this plasticity and adaptability is difficult, especially in fish that have longer lifespans. Finally, warming temperatures may not just affect spawning location directly but will have other effects consequences such as shifts spawning phenology, prey mismatch for larvae, and increased metabolic demand (Barbeaux et al., 2020; Hinckley et al., 2019; Rogers & Dougherty, 2019). These, and other factors, have likely affected and will continue affect the choice of spawning habitat in the future.

Predicting distributional shifts in response to warming – one of the ‘universal’ responses to climate change – is a central challenge as such shifts have far-reaching biological and socioeconomic implications (Pinsky, Selden, et al., 2020). Many studies rely on correlational relationships between environmental variables and biological variables such as abundance or biomass (e.g., Morley et al., 2018). Mechanistic-based approaches are gaining traction and improve upon the uncertainty in correlational predictions (Cote et al., 2021; Evans et al., 2015; Huey et al., 2012). For example, a species may not encounter the entire range of temperatures they can physiologically withstand (Kearney et al., 2020). Additionally, mechanistic-based approaches allow us to gain a deeper understanding of why certain distributional shifts may occur (Cheung et al., 2013; Deutsch et al., 2015, 2020). Finally, different life stages typically have different thermal sensitives, resulting in ontogenetic changes in distributions that are linked to habitat use (Barbeaux et al., 2020; Dahlke et al., 2020). Predicting distribution based on correlative approaches alone is often limiting in terms of what life stages can be observed (Fredston et al., 2021; Morley et al., 2018; Pinsky et al., 2013). While such approaches work for some life stages (juveniles, adults), they are difficult to extend to early life stages (eggs). Because early life stages may act as a bottleneck to a species’ ability to adapt to warming and other changing environmental conditions (Asch & Erisman, 2018; Ciannelli et al., 2021; Dahlke et al., 2018, 2020), understanding how warming will affect them is a critical step in understanding, predicting, and mitigating the effects of a changing climate. Here, we make use of a mechanistic relationship between hatch success and temperature and leverage global climate models that offer projections of temperature to understand how suitable spawning habitat – and thus the distribution of spawning adults, eggs, and newly-hatched larvae – has changed in the past and will continue to do so in the future. As polar regions like the Bering Sea are rapidly changing environmentally (Constable et al. 2022; Stabeno & Bell, 2019; Zhang, 2005), predictions such as ours will facilitate preparing for future uncertainty.

## Acknowledgements

LR acknowledges NPRB grant #. JSB acknowledges NSF grant number 2109411. We thank the Plankton Sorting and Identification Center (Poland) and Alaska Fisheries Science Center (USA) for their taxonomic expertise. Get ROMS team/ACLIM acks & grant numbers. EcoFOCI publication #.

## Data and code availability

All data and code necessary to reproduce the results in this study will be archived in github upon acceptance. We place no restrictions on data or code availability.

* probably need to add a clause about the CMIP6 projections not being publicly available

## References

Agostini, V. N., & Bakun, A. (2002). ‘Ocean triads’ in the Mediterranean Sea: Physical mechanisms potentially structuring reproductive habitat suitability (with example application to European anchovy, Engraulis encrasicolus): “Ocean triads” in the Mediterranean Sea. Fisheries Oceanography, 11(3), 129–142. https://doi.org/10.1046/j.1365-2419.2002.00201.x

Alderdice, D. F., & Forrester, C. R. (1971). Effects of Salinity, Temperature, and Dissolved Oxygen on Early Development of the Pacific Cod (Gadus macrocephalus). Journal of the Fisheries Research Board of Canada, 28(6), 883–902. https://doi.org/10.1139/f71-130

Asch, R. G., & Erisman, B. (2018). Spawning aggregations act as a bottleneck influencing climate change impacts on a critically endangered reef fish. Diversity and Distributions, 24(12), 1712–1728. https://doi.org/10.1111/ddi.12809

Barbeaux, S. J., & Hollowed, A. B. (2018). Ontogeny matters: Climate variability and effects on fish distribution in the eastern Bering Sea. Fisheries Oceanography, 27(1), 1–15. https://doi.org/10.1111/fog.12229

Barbeaux, S. J., Holsman, K., & Zador, S. (2020). Marine Heatwave Stress Test of Ecosystem-Based Fisheries Management in the Gulf of Alaska Pacific Cod Fishery. Frontiers in Marine Science, 7, 703. https://doi.org/10.3389/fmars.2020.00703

Baudron, A. R., Pecl, G., Gardner, C., Fernandes, P. G., & Audzijonyte, A. (2019). Ontogenetic deepening of Northeast Atlantic fish stocks is not driven by fishing exploitation. Proceedings of the National Academy of Sciences, 116(7), 2390–2392. https://doi.org/10.1073/pnas.1817295116

Beck, M. W., Heck, K. L., Able, K. W., Childers, D. L., Eggleston, D. B., Gillanders, B. M., Halpern, B., Hays, C. G., Hoshino, K., Minello, T. J., Orth, R. J., Sheridan, P. F., & Weinstein, M. P. (2001). The Identification, Conservation, and Management of Estuarine and Marine Nurseries for Fish and Invertebrates. BioScience, 51(8), 633. https://doi.org/10.1641/0006-3568(2001)051[0633:TICAMO]2.0.CO;2

Bennie, J., Hodgson, J. A., Lawson, C. R., Holloway, C. T. R., Roy, D. B., Brereton, T., Thomas, C. D., & Wilson, R. J. (2013). Range expansion through fragmented landscapes under a variable climate. Ecology Letters, 16(7), 921–929. https://doi.org/10.1111/ele.12129

Bian, X., Zhang, X., Sakurai, Y., Jin, X., Gao, T., Wan, R., & Yamamoto, J. (2014). Temperature-mediated survival, development and hatching variation of Pacific cod Gadus macrocephalus eggs: Temperature effects on g. macrocephalus eggs. Journal of Fish Biology, 84(1), 85–105. https://doi.org/10.1111/jfb.12257

Bian, X., Zhang, X., Sakurai, Y., Jin, X., Wan, R., Gao, T., & Yamamoto, J. (2016). Interactive effects of incubation temperature and salinity on the early life stages of pacific cod Gadus macrocephalus. Deep Sea Research Part II: Topical Studies in Oceanography, 124, 117–128. https://doi.org/10.1016/j.dsr2.2015.01.019

Blanchard, J. L., Maxwell, D. L., & Jennings, S. (2008). Power of monitoring surveys to detect abundance trends in depleted populations: The effects of density-dependent habitat use, patchiness, and climate change. ICES Journal of Marine Science, 65(1), 111–120. https://doi.org/10.1093/icesjms/fsm182

Brown, L. R., Komoroske, L. M., Wagner, R. W., Morgan-King, T., May, J. T., Connon, R. E., & Fangue, N. A. (2016). Coupled Downscaled Climate Models and Ecophysiological Metrics Forecast Habitat Compression for an Endangered Estuarine Fish. PLOS ONE, 11(1), e0146724. https://doi.org/10.1371/journal.pone.0146724

Bryan, D. R., McDermott, S. F., Nielsen, J. K., Fraser, D., & Rand, K. M. (2021). Seasonal migratory patterns of Pacific cod (Gadus macrocephalus) in the Aleutian Islands. Animal Biotelemetry, 9(1), 24. https://doi.org/10.1186/s40317-021-00250-2

Burrows, M. T., Schoeman, D. S., Richardson, A. J., Molinos, J. G., Hoffmann, A., Buckley, L. B., Moore, P. J., Brown, C. J., Bruno, J. F., Duarte, C. M., Halpern, B. S., Hoegh-Guldberg, O., Kappel, C. V., Kiessling, W., O’Connor, M. I., Pandolfi, J. M., Parmesan, C., Sydeman, W. J., Ferrier, S., … Poloczanska, E. S. (2014). Geographical limits to species-range shifts are suggested by climate velocity. Nature, 507(7493), 492–495. https://doi.org/10.1038/nature12976

Cheng, W., Hermann, A. J., Hollowed, A. B., Holsman, K. K., Kearney, K. A., Pilcher, D. J., Stock, C. A., & Aydin, K. Y. (2021). Eastern Bering Sea shelf environmental and lower trophic level responses to climate forcing: Results of dynamical downscaling from CMIP6. Deep Sea Research Part II: Topical Studies in Oceanography, 193, 104975. https://doi.org/10.1016/j.dsr2.2021.104975

Cheung, W. W. L., Lam, V. W. Y., Sarmiento, J. L., Kearney, K., Watson, R., & Pauly, D. (2009). Projecting global marine biodiversity impacts under climate change scenarios. Fish and Fisheries, 10(3), 235–251. https://doi.org/10.1111/j.1467-2979.2008.00315.x

Cheung, W. W. L., Sarmiento, J. L., Dunne, J., Frölicher, T. L., Lam, V. W. Y., Deng Palomares, M. L., Watson, R., & Pauly, D. (2013). Shrinking of fishes exacerbates impacts of global ocean changes on marine ecosystems. Nature Climate Change, 3(3), 254–258. https://doi.org/10.1038/nclimate1691

Ciannelli, L., Bailey, K., & Olsen, E. M. (2015). Evolutionary and ecological constraints of fish spawning habitats. ICES Journal of Marine Science, 72(2), 285–296. https://doi.org/10.1093/icesjms/fsu145

Ciannelli, L., Neuheimer, A. B., Stige, L. C., Frank, K. T., Durant, J. M., Hunsicker, M., Rogers, L. A., Porter, S., Ottersen, G., & Yaragina, N. A. (2021). Ontogenetic spatial constraints of sub-arctic marine fish species. Fish and Fisheries, 23(2), 342–357. https://doi.org/10.1111/faf.12619

Claydon, J. (2004). SPAWNING AGGREGATIONS OF CORAL REEF FISHES: CHARACTERISTICS, HYPOTHESES, THREATS AND MANAGEMENT. In R. N. Gibson, R. J. A. Atkinson, & J. D. M. Gordon (Eds.), Oceanography and Marine Biology (0 ed., pp. 273–310). CRC Press. https://doi.org/10.1201/9780203507810-11

Coachman, L. K. (1986). Circulation, water masses, and fluxes on the southeastern Bering Sea shelf. Continental Shelf Research, 5(1–2), 23–108. https://doi.org/10.1016/0278-4343(86)90011-7

Comte, L., & Olden, J. D. (2017). Climatic vulnerability of the world’s freshwater and marine fishes. Nature Climate Change, 7(10), 718–722. https://doi.org/10.1038/nclimate3382

Constable, A.J., S. Harper, J. Dawson, K. Holsman, T. Mustonen, D. Piepenburg, and B. Rost, 2022: Cross-Chapter Paper 6: Polar Regions. In: Climate Change 2022: Impacts, Adaptation and Vulnerability. Contribution of Working Group II to the Sixth Assessment Report of the Intergovernmental Panel on Climate Change [H.-O. Pörtner, D.C. Roberts, M. Tignor, E.S. Poloczanska, K. Mintenbeck, A. Alegría, M. Craig, S. Langsdorf, S. Löschke, V. Möller, A. Okem, B. Rama (eds.)]. Cambridge University Press, Cambridge, UK and New York, NY, USA, pp. 2319–2368, doi:10.1017/9781009325844.023.

Cote, D., Konecny, C. A., Seiden, J., Hauser, T., Kristiansen, T., & Laurel, B. J. (2021). Forecasted Shifts in Thermal Habitat for Cod Species in the Northwest Atlantic and Eastern Canadian Arctic. Frontiers in Marine Science, 8, 764072. https://doi.org/10.3389/fmars.2021.764072

Dahlke, F. T., Butzin, M., Nahrgang, J., Puvanendran, V., Mortensen, A., Pörtner, H.-O., & Storch, D. (2018). Northern cod species face spawning habitat losses if global warming exceeds 1.5°C. Science Advances, 4(11), eaas8821. https://doi.org/10.1126/sciadv.aas8821

Dahlke, F. T., Wohlrab, S., Butzin, M., & Pörtner, H.-O. (2020). Thermal bottlenecks in the life cycle define climate vulnerability of fish. Science, 369(6499), 65–70. https://doi.org/10.1126/science.aaz3658

Deutsch, C., Ferrel, A., Seibel, B., Pörtner, H.-O., & Huey, R. B. (2015). Climate change tightens a metabolic constraint on marine habitats. Science, 348(6239), 1132–1135. https://doi.org/10.1126/science.aaa1605

Deutsch, C., Penn, J. L., & Seibel, B. (2020). Metabolic trait diversity shapes marine biogeography. Nature, 585(7826), 557–562. https://doi.org/10.1038/s41586-020-2721-y

Dodson, J. J., Daigle, G., Hammer, C., Polte, P., Kotterba, P., Winkler, G., & Zimmermann, C. (2019). Environmental determinants of larval herring (Clupea harengus) abundance and distribution in the western Baltic Sea: Larval herring ecology. Limnology and Oceanography, 64(1), 317–329. https://doi.org/10.1002/lno.11042

Drost, H. E., Fisher, J., Randall, F., Kent, D., Carmack, E. C., & Farrell, A. P. (2016). Upper thermal limits of the hearts of Arctic cod Boreogadus saida: Adults compared with larvae: boreogadus saida thermal limits. Journal of Fish Biology, 88(2), 718–726. https://doi.org/10.1111/jfb.12807

Dulvy, N. K., Rogers, S. I., Jennings, S., Stelzenmller, V., Dye, S. R., & Skjoldal, H. R. (2008). Climate change and deepening of the North Sea fish assemblage: A biotic indicator of warming seas. Journal of Applied Ecology, 45(4), 1029–1039. https://doi.org/10.1111/j.1365-2664.2008.01488.x

Eaton, J. G., & Scheller, R. M. (1996). Effects of climate warming on fish thermal habitat in streams of the United States. Limnology and Oceanography, 41(5), 1109–1115. https://doi.org/10.4319/lo.1996.41.5.1109

Edwards, M., & Richardson, A. J. (2004). Impact of climate change on marine pelagic phenology and trophic mismatch. Nature, 430(7002), 881–884. https://doi.org/10.1038/nature02808

English, P. A., Ward, E. J., Rooper, C. N., Forrest, R. E., Rogers, L. A., Hunter, K. L., Edwards, A. M., Connors, B. M., & Anderson, S. C. (2021). Contrasting climate velocity impacts in warm and cool locations show that effects of marine warming are worse in already warmer temperate waters. Fish and Fisheries, 23(1), 239–255. https://doi.org/10.1111/faf.12613

Evans, T. G., Diamond, S. E., & Kelly, M. W. (2015). Mechanistic species distribution modelling as a link between physiology and conservation. Conservation Physiology, 3(1), cov056. https://doi.org/10.1093/conphys/cov056

Farley, E. V., Heintz, R. A., Andrews, A. G., & Hurst, T. P. (2016). Size, diet, and condition of age-0 Pacific cod (Gadus macrocephalus) during warm and cool climate states in the eastern Bering sea. Deep Sea Research Part II: Topical Studies in Oceanography, 134, 247–254. https://doi.org/10.1016/j.dsr2.2014.12.011

Fissel, B., Dalton, M, Garber-Yonts, B, Haynie, A. C., Kasperski, S., Lee, Jean, Lew, D, Lavoie, A, Seung, Chang, Sparks, K, Szymkowiak, Marysia, & Wise, S. (2019). Stock Assessment and Fishery Evaluation Report for the groundfish fisheries of the Gulf of Alaska and Bering Sea/Aleutian Islands Area: Economic Status of the Groundfish Fisheries Off Alaska (p. 385) [Groundfish SAFE Economic Report].

Fossheim, M., Primicerio, R., Johannesen, E., Ingvaldsen, R. B., Aschan, M. M., & Dolgov, A. V. (2015). Recent warming leads to a rapid borealization of fish communities in the Arctic. Nature Climate Change, 5(7), 673–677. https://doi.org/10.1038/nclimate2647

Fredston, A., Pinsky, M., Selden, R. L., Szuwalski, C., Thorson, J. T., Gaines, S. D., & Halpern, B. S. (2021). Range edges of North American marine species are tracking temperature over decades. Global Change Biology, 27(13), 3145–3156. https://doi.org/10.1111/gcb.15614

García Molinos, J., Halpern, B. S., Schoeman, D. S., Brown, C. J., Kiessling, W., Moore, P. J., Pandolfi, J. M., Poloczanska, E. S., Richardson, A. J., & Burrows, M. T. (2016). Climate velocity and the future global redistribution of marine biodiversity. Nature Climate Change, 6(1), 83–88. https://doi.org/10.1038/nclimate2769

Grebmeier, J. M. (2006). A Major Ecosystem Shift in the Northern Bering Sea. Science, 311(5766), 1461–1464. https://doi.org/10.1126/science.1121365

Hawkins, E., Osborne, T. M., Ho, C. K., & Challinor, A. J. (2013). Calibration and bias correction of climate projections for crop modelling: An idealised case study over Europe. Agricultural and Forest Meteorology, 170, 19–31. https://doi.org/10.1016/j.agrformet.2012.04.007

Hayes, D. B., Ferreri, C. P., & Taylor, W. W. (1996). Linking fish habitat to their population dynamics. 53, 8.

Haynie, A. C., & Pfeiffer, L. (2013). Climatic and economic drivers of the Bering Sea walleye pollock (Theragra chalcogramma) fishery: Implications for the future. Canadian Journal of Fisheries and Aquatic Sciences, 70(6), 841–853. https://doi.org/10.1139/cjfas-2012-0265

Hermann, A. J., Kearney, K., Cheng, W., Pilcher, D., Aydin, K., Holsman, K. K., & Hollowed, A. B. (2021). Coupled modes of projected regional change in the Bering Sea from a dynamically downscaling model under CMIP6 forcing. Deep Sea Research Part II: Topical Studies in Oceanography, 194, 104974. https://doi.org/10.1016/j.dsr2.2021.104974

Hinckley, S., Stockhausen, W. T., Coyle, K. O., Laurel, B. J., Gibson, G. A., Parada, C., Hermann, A. J., Doyle, M. J., Hurst, T. P., Punt, A. E., & Ladd, C. (2019). Connectivity between spawning and nursery areas for Pacific cod (Gadus macrocephalus) in the Gulf of Alaska. Deep Sea Research Part II: Topical Studies in Oceanography, 165, 113–126. https://doi.org/10.1016/j.dsr2.2019.05.007

Hollowed, A. B., Hare, S. R., & Wooster, W. S. (2001). Pacific Basin climate variability and patterns of Northeast Pacific marine fish production. Progress in Oceanography, 49(1–4), 257–282. https://doi.org/10.1016/S0079-6611(01)00026-X

Holsman, K. K., Haynie, A. C., Hollowed, A. B., Reum, J. C. P., Aydin, K., Hermann, A. J., Cheng, W., Faig, A., Ianelli, J. N., Kearney, K. A., & Punt, A. E. (2020). Ecosystem-based fisheries management forestalls climate-driven collapse. Nature Communications, 11(1), 4579. https://doi.org/10.1038/s41467-020-18300-3

Houde, E. D. (2008). Emerging from Hjort’s Shadow. Journal of Northwest Atlantic Fishery Science, 41, 53–70. https://doi.org/10.2960/J.v41.m634

Huey, R. B., Kearney, M. R., Krockenberger, A., Holtum, J. A. M., Jess, M., & Williams, S. E. (2012). Predicting organismal vulnerability to climate warming: Roles of behaviour, physiology and adaptation. Philosophical Transactions of the Royal Society B: Biological Sciences, 367(1596), 1665–1679. https://doi.org/10.1098/rstb.2012.0005

Hurst, T. P., Munch, S. B., & Lavelle, K. A. (2012). Thermal reaction norms for growth vary among cohorts of Pacific cod (Gadus macrocephalus). Marine Biology, 159(10), 2173– 2183. https://doi.org/10.1007/s00227-012-2003-9

Kearney, K., Hermann, A., Cheng, W., Ortiz, I., & Aydin, K. (2020). A coupled pelagic–benthic– sympagic biogeochemical model for the Bering Sea: Documentation and validation of the BESTNPZ model (v2019.08.23) within a high-resolution regional ocean model. Geoscientific Model Development, 13(2), 597–650. https://doi.org/10.5194/gmd-13-597-2020

Kleisner, K. M., Fogarty, M. J., McGee, S., Hare, J. A., Moret, S., Perretti, C. T., & Saba, V. S. (2017). Marine species distribution shifts on the U.S. Northeast Continental Shelf under continued ocean warming. Progress in Oceanography, 153, 24–36. https://doi.org/10.1016/j.pocean.2017.04.001

Kotwicki, S., & Lauth, R. R. (2013). Detecting temporal trends and environmentally-driven changes in the spatial distribution of bottom fishes and crabs on the eastern Bering Sea shelf. Deep Sea Research Part II: Topical Studies in Oceanography, 94, 231–243. https://doi.org/10.1016/j.dsr2.2013.03.017

Laurel, B. J., Hurst, T. P., Copeman, L. A., & Davis, M. W. (2008). The role of temperature on the growth and survival of early and late hatching Pacific cod larvae (Gadus macrocephalus). Journal of Plankton Research, 30(9), 1051–1060. https://doi.org/10.1093/plankt/fbn057

Laurel, B. J., & Rogers, L. A. (2020). Loss of spawning habitat and prerecruits of Pacific cod during a Gulf of Alaska heatwave. Canadian Journal of Fisheries and Aquatic Sciences, 77(4), 644–650. https://doi.org/10.1139/cjfas-2019-0238

Litzow, M. A., Abookire, A. A., Duffy-Anderson, J. T., Laurel, B. J., Malick, M. J., & Rogers, L. A. (2022). Predicting year class strength for climate-stressed gadid stocks in the Gulf of Alaska. Fisheries Research, 249, 106250. https://doi.org/10.1016/j.fishres.2022.106250

Marsh, J. M., & Mueter, F. J. (2020). Influences of temperature, predators, and competitors on polar cod (Boreogadus saida) at the southern margin of their distribution. Polar Biology, 43(8), 995–1014. https://doi.org/10.1007/s00300-019-02575-4

Matarese, A. C., Blood, D. M., Picquelle, S. J., & Benson, J. L. (2003). Atlas of Abundance and Distribution Patterns of Ichthyoplankton from the Northeast Pacific Ocean and Bering Sea Ecosystems Based on Research Conducted by the Alaska Fisheries Science Center (1972-1996).

McKenzie, D. J., Zhang, Y., Eliason, E. J., Schulte, P. M., Claireaux, G., Blasco, F. R., Nati, J. J. H., & Farrell, A. P. (2020). Intraspecific variation in tolerance of warming in fishes. Journal of Fish Biology, 98(6), 1536–1555. https://doi.org/10.1111/jfb.14620

Miller, J. A., DiMaria, R. A., & Hurst, T. P. (2016). Patterns of larval source distribution and mixing in early life stages of Pacific cod (Gadus macrocephalus) in the southeastern Bering Sea. Deep Sea Research Part II: Topical Studies in Oceanography, 134, 270– 282. https://doi.org/10.1016/j.dsr2.2014.12.012

Mizobata, K., Wang, J., & Saitoh, S. (2006). Eddy-induced cross-slope exchange maintaining summer high productivity of the Bering Sea shelf break. Journal of Geophysical Research, 111(C10), C10017. https://doi.org/10.1029/2005JC003335

Morley, J. W., Selden, R. L., Latour, R. J., Frölicher, T. L., Seagraves, R. J., & Pinsky, M. L. (2018). Projecting shifts in thermal habitat for 686 species on the North American continental shelf. PLOS ONE, 13(5), e0196127. https://doi.org/10.1371/journal.pone.0196127

Mueter, F. J., & Litzow, M. A. (2008). SEA ICE RETREAT ALTERS THE BIOGEOGRAPHY OF THE BERING SEA CONTINENTAL SHELF. Ecological Applications, 18(2), 309–320. https://doi.org/10.1890/07-0564.1

Munday, P. L., Jones, G. P., Pratchett, M. S., & Williams, A. J. (2008). Climate change and the future for coral reef fishes. Fish and Fisheries, 9(3), 261–285. https://doi.org/10.1111/j.1467-2979.2008.00281.x

Neidetcher, S. K., Hurst, T. P., Ciannelli, L., & Logerwell, E. A. (2014). Spawning phenology and geography of Aleutian Islands and eastern Bering Sea Pacific cod (Gadus macrocephalus). Deep Sea Research Part II: Topical Studies in Oceanography, 109, 204–214. https://doi.org/10.1016/j.dsr2.2013.12.006

Overland, J. E., Wang, M., Wood, K. R., Percival, D. B., & Bond, N. A. (2012). Recent Bering Sea warm and cold events in a 95-year context. Deep Sea Research Part II: Topical Studies in Oceanography, 65–70, 6–13. https://doi.org/10.1016/j.dsr2.2012.02.013

Parmesan, C., & Yohe, G. (2003). A globally coherent fingerprint of climate change impacts across natural systems. Nature, 421(6918), 37–42. https://doi.org/10.1038/nature01286

Perry, A. L., Low, P. J., Ellis, J. R., & Reynolds, J. D. (2005). Climate Change and Distribution Shifts in Marine Fishes. Science, 308(5730), 1912–1915. https://doi.org/10.1126/science.1111322

Pilcher, D.J., Naiman, D.M., Cross, J.N., Hermann, A.J., Siedlecki, S.A., Gibson, G.A. and Mathis, J.T., (2019). Modeled effect of coastal biogeochemical processes, climate variability, and ocean acidification on aragonite saturation state in the Bering Sea. Frontiers in Marine Science, 5:508; https://doi.org/10.3389/fmars.2018.00508

Pinsky, M. L., & Fogarty, M. (2012). Lagged social-ecological responses to climate and range shifts in fisheries. Climatic Change, 115(3–4), 883–891. https://doi.org/10.1007/s10584-012-0599-x

Pinsky, M. L., Rogers, L. A., Morley, J. W., & Frölicher, T. L. (2020). Ocean planning for species on the move provides substantial benefits and requires few trade-offs. Science Advances, 6(50). https://doi.org/10.1126/sciadv.abb8428

Pinsky, M. L., Selden, R. L., & Kitchel, Z. J. (2020). Climate-Driven Shifts in Marine Species Ranges: Scaling from Organisms to Communities. Annual Review of Marine Science, 12(1), 153–179. https://doi.org/10.1146/annurev-marine-010419-010916

Pinsky, M. L., Worm, B., Fogarty, M. J., Sarmiento, J. L., & Levin, S. A. (2013). Marine Taxa Track Local Climate Velocities. Science, 341(6151), 1239–1242. https://doi.org/10.1126/science.1239352

Poloczanska, E. S., Brown, C. J., Sydeman, W. J., Kiessling, W., Schoeman, D. S., Moore, P. J., Brander, K., Bruno, J. F., Buckley, L. B., Burrows, M. T., Duarte, C. M., Halpern, B. S., Holding, J., Kappel, C. V., O’Connor, M. I., Pandolfi, J. M., Parmesan, C., Schwing, F., Thompson, S. A., & Richardson, A. J. (2013). Global imprint of climate change on marine life. Nature Climate Change, 3(10), 919–925. https://doi.org/10.1038/nclimate1958

Porter, S. M., & Ciannelli, L. (2018). Effect of temperature on Flathead Sole (Hippoglossoides elassodon) spawning in the southeastern Bering Sea during warm and cold years. Journal of Sea Research, 141, 26–36. https://doi.org/10.1016/j.seares.2018.08.003

Pörtner, H. O., & Farrell, A. P. (2008). Physiology and Climate Change. Science, 322(5902), 690–692. https://doi.org/10.1126/science.1163156

Pörtner, H.-O. (2021). Climate impacts on organisms, ecosystems and human societies: Integrating OCLTT into a wider context. Journal of Experimental Biology, 224(Suppl_1), jeb238360. https://doi.org/10.1242/jeb.238360

Rand, K. M., Munro, P., Neidetcher, S. K., & Nichol, D. G. (2014). Observations of Seasonal Movement from a Single Tag Release Group of Pacific Cod in the Eastern Bering Sea. Marine and Coastal Fisheries, 6(1), 287–296. https://doi.org/10.1080/19425120.2014.976680

Rijnsdorp, A. D., Peck, M. A., Engelhard, G. H., Möllmann, C., & Pinnegar, J. K. (2009). Resolving the effect of climate change on fish populations. ICES Journal of Marine Science, 66(7), 1570–1583. https://doi.org/10.1093/icesjms/fsp056

Rogers, L. A., & Dougherty, A. B. (2019). Effects of climate and demography on reproductive phenology of a harvested marine fish population. Global Change Biology, 25(2), 708– 720. https://doi.org/10.1111/gcb.14483

Rogers, L. A., Griffin, R., Young, T., Fuller, E., St. Martin, K., & Pinsky, M. L. (2019). Shifting habitats expose fishing communities to risk under climate change. Nature Climate Change, 9(7), 512–516. https://doi.org/10.1038/s41558-019-0503-z

Rogers, L. A., Wilson, M. T., Duffy-Anderson, J. T., Kimmel, D. G., & Lamb, J. F. (2021). Pollock and “the Blob”: Impacts of a marine heatwave on walleye pollock early life stages. Fisheries Oceanography, 30(2), 142–158. https://doi.org/10.1111/fog.12508

Saha, S., Nadiga, S., Thiaw, C., Wang, J., Wang, W., Zhang, Q., Van den Dool, H.M., Pan, H.L., Moorthi, S., Behringer, D. and Stokes, D., (2006). The NCEP climate forecast system. Journal of Climate, 19(15), pp.3483–3517.

Sample, T., & Wolotira, Jr, R. (1985). Demersal Fish and Shellfish Resources of Norton Sound and Adjacent Waters During 1979. NOAA Technical Memorandum, NMFS F/NWC-89.

Shimada, A. M., & Kimura, D. K. (1994). Seasonal movements of Pacific cod, Gadus macrocephalUS, in the eastern Bering Sea and adjacent waters based on tag-recapture data. FISHERY BULLETIN, 92, 800–816.

Sigler, M. F., Harvey, H. R., Ashjian, J., Lomas, M. W., Napp, J. M., Stabeno, P. J., & Van Pelt, T. I. (2010). How Does Climate Change Affect the Bering Sea Ecosystem? Eos, Transactions American Geophysical Union, 91(48), 457. https://doi.org/10.1029/2010EO480001

Smart, T., Duffy-Anderson, J., & Horne, J. (2012). Alternating temperature states influence walleye pollock early life stages in the southeastern Bering Sea. Marine Ecology Progress Series, 455, 257–267. https://doi.org/10.3354/meps09619

Spies, I., Gruenthal, K. M., Drinan, D. P., Hollowed, A. B., Stevenson, D. E., Tarpey, C. M., & Hauser, L. (2019). Genetic evidence of a northward range expansion in the eastern Bering Sea stock of Pacific cod. Evolutionary Applications, 13(2), 362–375. https://doi.org/10.1111/eva.12874

Springer, V. G. (1961). Notes on and Additions to the Fish Fauna of the Tampa Bay Area in Florida. Copeia, 1961(4), 480. https://doi.org/10.2307/1439600

Stabeno, P. J., & Bell, S. W. (2019). Extreme Conditions in the Bering Sea (2017–2018): Record-Breaking Low Sea-Ice Extent. Geophysical Research Letters, 46(15), 8952– 8959. https://doi.org/10.1029/2019GL083816

Stabeno, P. J., Bond, N. A., Kachel, N. B., Salo, S. A., & Schumacher, J. D. (2001). On the temporal variability of the physical environment over the south-eastern Bering Sea: Bering Sea physical conditions. Fisheries Oceanography, 10(1), 81–98. https://doi.org/10.1046/j.1365-2419.2001.00157.x

Stabeno, P. J., Danielson, S. L., Kachel, D. G., Kachel, N. B., & Mordy, C. W. (2016). Currents and transport on the Eastern Bering Sea shelf: An integration of over 20 years of data. Deep Sea Research Part II: Topical Studies in Oceanography, 134, 13–29. https://doi.org/10.1016/j.dsr2.2016.05.010

Stabeno, P. J., Duffy-Anderson, J. T., Eisner, L. B., Farley, E. V., Heintz, R. A., & Mordy, C. W. (2017). Return of warm conditions in the southeastern Bering Sea: Physics to fluorescence. PLOS ONE, 12(9), e0185464. https://doi.org/10.1371/journal.pone.0185464

Stabeno, P. J., Farley Jr., E. V., Kachel, N. B., Moore, S., Mordy, C. W., Napp, J. M., Overland, J. E., Pinchuk, A. I., & Sigler, M. F. (2012). A comparison of the physics of the northern and southern shelves of the eastern Bering Sea and some implications for the ecosystem. Deep Sea Research Part II: Topical Studies in Oceanography, 65–70, 14– 30. https://doi.org/10.1016/j.dsr2.2012.02.019

Stabeno, P. J., Kachel, N. B., Moore, S. E., Napp, J. M., Sigler, M., Yamaguchi, A., & Zerbini, A. N. (2012). Comparison of warm and cold years on the southeastern Bering Sea shelf and some implications for the ecosystem. Deep Sea Research Part II: Topical Studies in Oceanography, 65–70, 31–45. https://doi.org/10.1016/j.dsr2.2012.02.020

Stark, J. (2007). Geographic and seasonal variations in maturation and growth pf female Pacific cod (Gadus macrocephalus) in the Gulf of Alaska and Bering Sea. FISHERY BULLETIN, 105, 396–407.

Stevenson, D. E., & Lauth, R. R. (2019). Bottom trawl surveys in the northern Bering Sea indicate recent shifts in the distribution of marine species. Polar Biology, 42(2), 407–421. https://doi.org/10.1007/s00300-018-2431-1

Sumaila, U. R., Cheung, W. W. L., Lam, V. W. Y., Pauly, D., & Herrick, S. (2011). Climate change impacts on the biophysics and economics of world fisheries. Nature Climate Change, 1(9), 449–456. https://doi.org/10.1038/nclimate1301

Sunday, J. M., Bates, A. E., & Dulvy, N. K. (2011). Global analysis of thermal tolerance and latitude in ectotherms. Proceedings of the Royal Society B: Biological Sciences, 278(1713), 1823–1830. https://doi.org/10.1098/rspb.2010.1295

Thompson, G. G., Barbeaux, S., Conner, J., Fissel, B., Hurst, T., Laurel, B., O’Leary, C. A., Rogers, L., Shotwell, S. K., Siddon, E., Spies, I., Thorson, J. T., & Tyrell, A. (2021). 2. Assessment of the Pacific Cod Stock in the Eastern Bering Sea. 494.

Wolotira, Jr, R., Sample, T., & Morin, Martin. (1977). Demersal Fish and Shellfish Resources of Norton Sounds, the southeastern Chukchi Sea, and adjacent waters in the baseline year 1976. Northwest and Alaska FIsheries Center Processed Report.

Zhang, J. (2005). Warming of the arctic ice-ocean system is faster than the global average since the 1960s: FASTER WARMING ARCTIC ICE-OCEAN SYSTEM. Geophysical Research Letters, 32(19), n/a-n/a. https://doi.org/10.1029/2005GL024216

