## Supplementary material for "Predicting Pacific cod spawning habitat in a changing climate": SI

This file contains

Figures S1 – S3

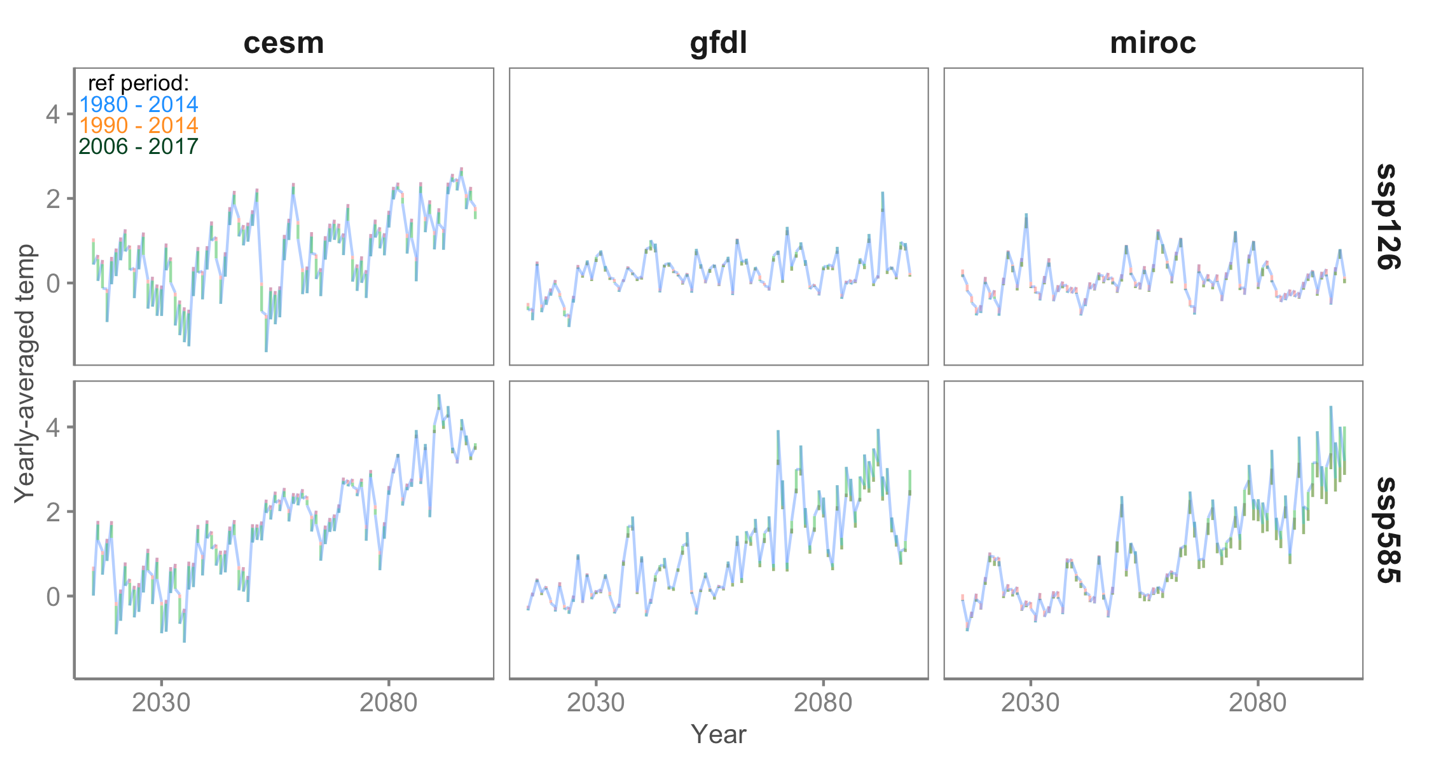

Figure S1. The bottom temperature projections for all models (columns) and emission scenarios (rows) are robust to choice of reference period for bias correction (blue, 1980 – 2014; orange, 1990 – 2014, and black, 2006 – 2017).

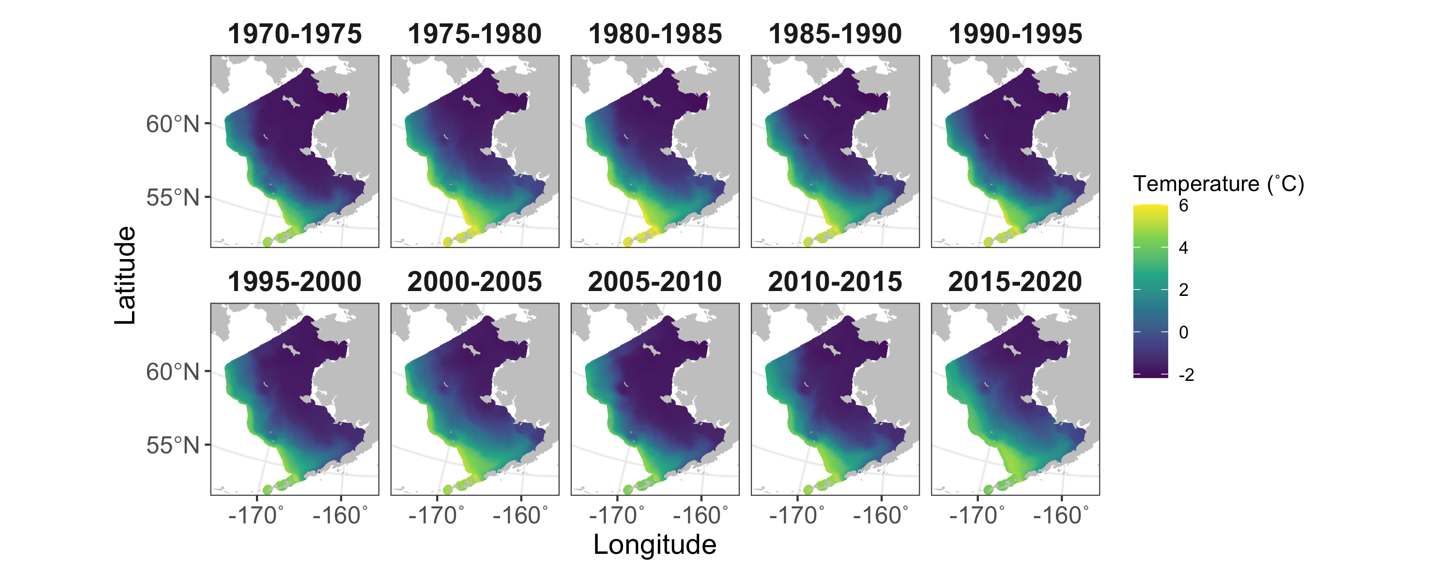

Figure S2. Bottom temperatures averaged over January – April across 5-year periods during the hindcast period (1970 – 2020).

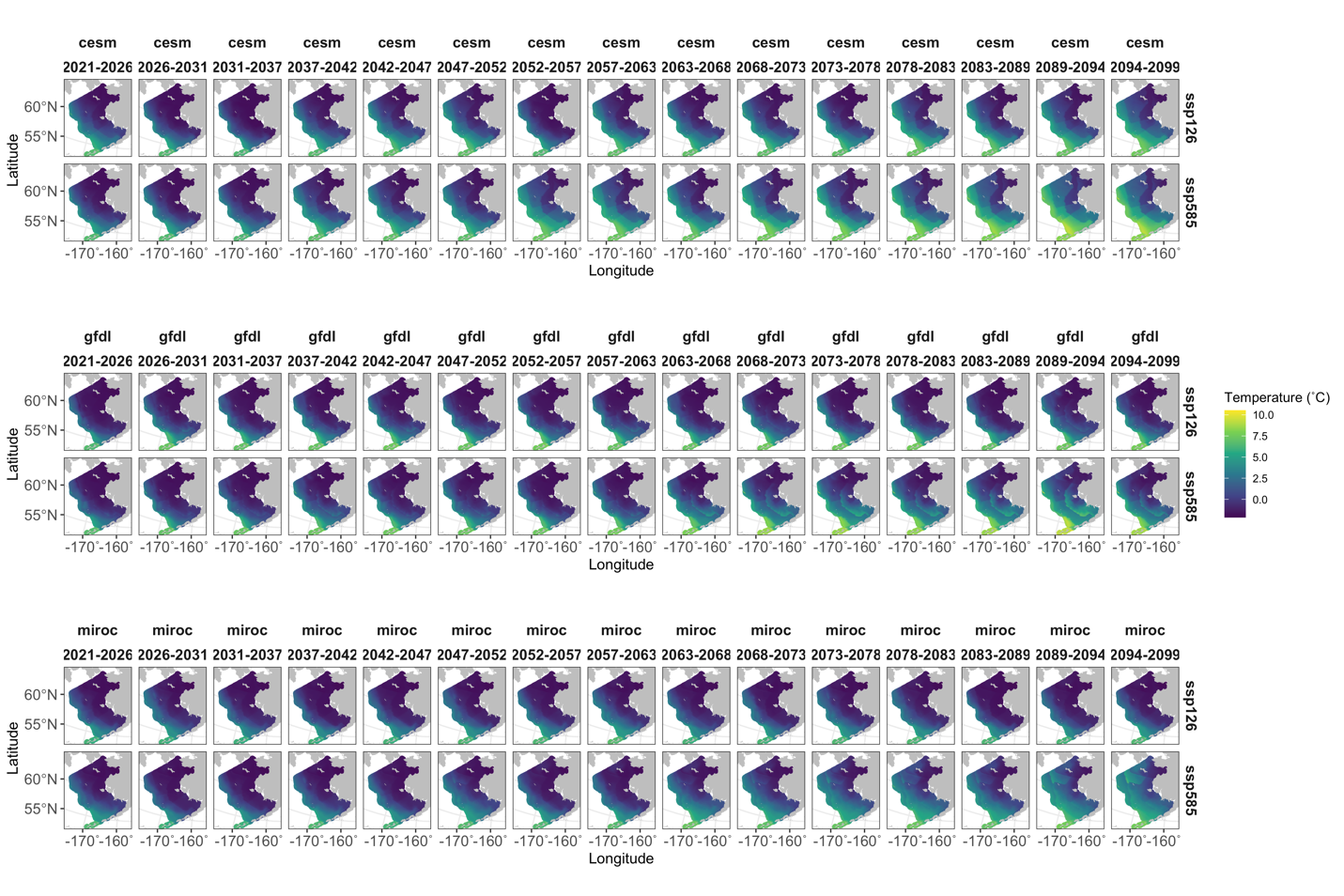

Figure S3. Bottom temperatures averaged over January – April across 5-year periods during the projection period (2021 - 2099) for all three global climate models downscaled for the Bering Sea in Bering 10k ROMS (top: CESM, middle: GFDL, bottom: MIROC).
